# Structural Degeneration of the Nucleus basalis of Meynert in Mild Cognitive Impairment and Alzheimer’s Disease – Evidence from an MRI-based Meta-Analysis

**DOI:** 10.1101/2023.09.06.556501

**Authors:** Marthe Mieling, Hannah Meier, Nico Bunzeck

## Abstract

Recent models of Alzheimer’s Disease (AD) suggest that neuropathological changes of the medial temporal lobe, especially entorhinal cortex, are preceded by degenerations of the cholinergic Nucleus basalis of Meynert (NbM). Evidence from imaging studies in humans, however, is limited. Therefore, we performed an activation-likelihood estimation meta-analysis on whole brain voxel-based morphometry (VBM) MRI data from 54 experiments and 2581 subjects in total. It revealed, compared to healthy older controls, reduced gray matter in the bilateral NbM in AD, but only limited evidence for such an effect in patients with mild cognitive impairment (MCI), which typically precedes AD. Both patient groups showed less gray matter in the amygdala and hippocampus, with hints towards more pronounced amygdala effects in AD. We discuss our findings in the context of studies that highlight the importance of the cholinergic basal forebrain in learning and memory throughout the life span, and conclude that they are partly compatible with pathological staging models suggesting initial and pronounced structural degenerations within the NbM in the progression of AD.

## Introduction

### Anatomy and function of the cholinergic basal forebrain

The basal forebrain (BF) cholinergic system is a subcortical structure located in the frontal cortex. It can be subdivided into four nuclei with predominantly cholinergic neurons, namely, the medial septal nucleus (Ch1), the vertical and horizontal limb of the diagonal band of Broca nuclei (Ch2, Ch3), and the magnocellular complex (Ch4), which mainly includes the Nucleus basalis of Meynert (NbM) (Mesulam et al., 1983). The NbM is considered to be the largest of the BF’s nuclei with ca. 13-14 mm from anterior-posterior and 16-18 mm from medial-lateral (Mesulam and Geula, 1988), and has, therefore, been targeted both in animal and human imaging studies. Functionally, the NbM plays an essential role in the modulation of complex behaviors and cognition through the communication with limbic structures and the entire neocortex (Mesulam et al., 1983), which we further explain below. Finally, cholinergic neurons within the NbM are complexly branched and, important for the understanding of life-long development, the axonal arborizations and synapses are required to remodel continuously due to their principal role in learning, memory and attention (Hasselmo, 2006; Mitsushima et al., 2013; Schmitz and Duncan, 2018; Wu et al., 2014).

Acetylcholine (ACh) has long been identified as a powerful neuromodulator that is involved in the regulation of neural activity in distant brain regions and thereby serves several functions, including learning and memory (Picciotto et al., 2012). For instance, the processing of novel stimuli is associated with neural responses in the monkey cholinergic BF (Wilson and Rolls, 1990) as well as increases in fronto-cortical and hippocampal acetylcholine levels in rats (Acquas et al., 1996). Further, high levels of ACh in the rat perirhinal cortex during the encoding of novel information promote memory performance, but they have a detrimental effect on consolidation (Winters et al., 2006) (see Gais and Born, 2004, for a similar effect in humans). Compatible with this observation, the removal of cholinergic inputs to the perirhinal cortex impairs object recognition in rodents (Winters and Bussey, 2005).

In humans, pharmacological stimulation of cholinergic activity by acetylcholinesterase inhibition leads to a change in neural novelty signals within the medial temporal lobe (MTL), including the parahippocampal cortex and hippocampus (Bunzeck et al., 2014). In electrophysiological studies, also in humans, cholinergic stimulation led to a shift in novelty responsive brain regions from medio-temporal to prefrontal areas, which suggests that the influence of the MTL and prefrontal regions in novelty processing is mediated by acetylcholine levels (Eckart and Bunzeck, 2013). Indeed, cholinergic antagonists can impair working memory (Aigner and Mishkin, 1986) and the encoding of novel information in explicit memory tasks (Sherman et al., 2003). In contrast, cholinergic agonists can have opposite effects (Buccafusco et al., 2005). Finally, acetylcholine also modulates attentional processing (Bauer et al., 2012; Hasselmo and Sarter, 2011) as well as human working memory and subsequent familiarity based recognition (Eckart et al., 2016) via changes in neural oscillations.

While the involvement of ACh in cognition, especially learning and memory, is widely accepted, the underlying mechanisms are still under debate. In this regard several effects of ACh within the MTL seem particularly important (Hasselmo, 2006). For instance, ACh enhances the afferent input from dentate gyrus and entorhinal cortex to CA3 (Giocomo and Hasselmo, 2005; Radcliffe et al., 1999); ACh is closely related to hippocampal theta rhythm (Bland and Oddie, 2001; Siok et al., 2006), which was linked to mnemonic processes (Herweg 2020); ACh can increase synaptic plasticity within hippocampal CA1 region (Adams et al., 2004; Huerta and Lisman, 1993) and entorhinal cortex (Cheong et al., 2001); and finally, ACh can enhance encoding related spike activity for novel information in the entorhinal cortex (Klink and Alonso, 1997). Computational models suggest (Gluck et al., 2005; Myers et al., 1996) that cholinergic projections from the BF, in particular the medial septum, to the hippocampus influence suppression of synaptic transmission within the hippocampus and the neocortex. This suppression is selective in as much it affects intrinsic, recurrent collaterals (such as CA1 and CA3) more strongly than external afferents (such as from the entorhinal cortex to CA1) of the hippocampal formation (Hasselmo et al., 1995; Myers et al., 1996).

Together, the basal forebrain, including the NbM, is a major source of cholinergic projections to the MTL and neocortex. Thereby, it has the potential to promote cognitive functions, including learning and memory, via the modulation of spike activity, neural oscillations and network properties (Picciotto et al., 2012; Záborszky et al., 2018).

### Age related changes of the basal forebrain

Structural degenerations of the BF can be observed during healthy aging, which leads to cognitive impairment (Düzel et al., 2010; Heys et al., 2010; Mesulam, 2004). While this can be distinguished from more prominent pathological changes in mild cognitive impairment (MCI) and Alzheimer’s disease (AD), it is also clear that healthy and pathological aging can be described on a continuum and, therefore, understanding developmental trajectories is essential. For instance, aged rats showed significant atrophy and a loss of cholinergic BF neurons (de Lacalle et al., 1996), which, in another study, closely related to specific cognitive impairments, including spatial learning (Fischer et al., 1989). Further, magnocellular BF cholinergic neurons are vulnerable to intraneuronal amyloid-β accumulation even in cognitively unimpaired healthy older humans, as shown in a post mortem study (Baker-Nigh et al., 2015).

With regard to disease progression, MCI is typically defined by mild cognitive problems, including memory and language, that are more pronounced than healthy age-related changes, but only minimally interfere with daily life. People with MCI are at higher risk of developing AD, which is characterized by even more pronounced cognitive impairments and structural degenerations (Jessen et al., 2014; Petersen et al., 2018). Physiologically, the prevailing view on the development of AD is that deposits of amyloid-β and p-tau first occur in the trans-entorhinal and entorhinal cortex (EC), located within the MTL, followed by the hippocampus and other cortical regions (Braak and Braak, 1991; Corder et al., 2000; Duyckaerts et al., 1990). Therefore, previous MRI studies in humans have often focused on structural abnormalities of these brain regions using voxel-based morphometry (VBM), or similar measures (Ashburner and Friston, 2000), and more recently, positron emission tomography (PET), that allows to quantify amyloid-β and pTau *in vivo* (Jagust, 2018). They could show that the volume (Karas et al., 2004) and shape (Gerardin et al., 2009) of the hippocampus as well as hippocampal subfields (Hett et al., 2019) differ between MCI and AD patients as compared to age-matched healthy controls (for a review see Frisoni et al., 2010). Similar but more heterogeneous effects have been observed in the EC, amygdala, and frontal and parietal cortex (Arrondo et al., 2022a).

The focus on MTL regions as origin of AD has been challenged by histologic studies (Geula and Mesulam, 1996; Mesulam et al., 2004; Schliebs and Arendt, 2011) and *in vivo* imaging that shows early pathological changes in the NbM (Grothe et al., 2013, 2012). Specifically, the BF was suggested as one of the first sub-cortical regions affected by neuronal loss (Davies and Maloney, 1976; Whitehouse et al., 1981), possibly due to accumulation of amyloid-ß and pTau, which is followed by decreased levels of ACh (Rajmohan and Reddy, 2017). This, in turn, could interfere with neuronal signaling, especially in memory encoding (Atri et al., 2004) and attentional processing (Ballinger et al., 2016). Therefore, major damages of cholinergic neurons of the BF in AD not only leads to a 90-95% loss in cortical ACh activity but also cognitive and behavioral impairments (Ballinger et al., 2016; Geula et al., 2021). Finally, in advanced AD neural loss is more pronounced in the NbM as compared to layer-II EC (Arendt et al., 2015).

Based on these findings, a recent view suggests the NbM as early origin of structural degeneration followed by the EC and other cortical brain regions. In humans, direct evidence comes from anatomical MRI studies in combination with measures of amyloid-β and pTau (Fernández-Cabello et al., 2020; Schmitz and Spreng, 2016). They revealed more pronounced NbM gray matter loss in cognitively healthy humans depending on abnormal vs normal CSF biomarkers (Fernández-Cabello et al., 2020; Schmitz et al., 2020; Schmitz and Spreng, 2016). Importantly, NbM baseline volumes predicted longitudinal EC degenerations, implying a trans-synaptic spread of amyloid-β starting in the NbM (Fernández-Cabello et al., 2020; Schmitz and Spreng, 2016). This conclusion is compatible with work in animals and adds a crucial upstream link to the subsequent spread from EC to other MTL structures, including the hippocampus, and more distant neocortical brain regions such as the posterior parietal cortex (de Calignon et al., 2012; Khan et al., 2014; Liu et al., 2012; Wu et al., 2016). Apart from these and other studies (e.g., Arrondo et al., 2022; Colloby et al., 2014; Fernández-Cabello et al., 2020; Jessen et al., 2006; Kerbler et al., 2015; Kilimann et al., 2014; Schmitz et al., 2018; Schmitz and Spreng, 2016) showing structural degenerations of the BF in vivo with MRI, others have demonstrated reduced structural integrity of cholinergic white matter pathways in AD (Schumacher et al. 2022, Nemy et al., 2023).

### Research Question and hypotheses

Despite the prominent view of a role of the cholinergic BF in AD, empirical evidence is limited, especially *in vivo*, and a meta-analytic investigation is missing. Here, we assumed that AD and MCI are characterized by smaller NbM volumes as compared to healthy controls (HC). More specifically, we hypothesized that patients with MCI show less gray matter in the NbM compared to age matched HC (HC>MCI, hypothesis 1); patients with AD show less gray matter in the NbM compared to age matched HC (HC>AD, hypothesis 2); and patients with MCI or AD show less gray matter in the MTL (EC and hippocampus, hypothesis 3) compared to age matched HC (HC>MCI/AD). To this end, we employed an activation likelihood estimation (ALE) meta-analytic approach on the basis of recently published whole brain voxel-based morphometry (VBM) MRI data. This allowed us to focus on the NbM and EC, and, in a more exploratory fashion, other brain regions, such as temporoparietal areas, which have also been associated with AD (Fernández-Cabello et al., 2020).

## Methods

### Literature search and selection

This work followed the PRISMA 2020 guidelines (Page et al., 2021) and is in accordance with suggestions by Müller et al. (2018) for Neuroimaging Meta-Analyses (see Table S1). More specifically, a systematic review was conducted based on the VBM BrainMap database using the search application Sleuth (version 3.0.4) (Fox et al., 2005; Fox and Lancaster, 2002; Vanasse et al., 2018). Here, the rationale is to identify those studies with convergence regarding brain volume reductions in MCI and AD compared to HC; therefore, literature review and statistical analyses (see below) were run separately for HC>MCI and HC>AD (Eickhoff et al., 2009; Vanasse et al., 2018).

The search included all whole-brain VBM studies in the database until Aug. 16, 2022 – these were 1316 publications with 4345 experiments (Fox et al., 2005; Fox and Lancaster, 2002; Vanasse et al., 2018). Here, the term experiment refers to a single analysis conducted in a given publication (Turkeltaub et al., 2012). Applying a specific category selection in Sleuth (see Table S1) and specific inclusion and exclusion criteria (Table 1) resulted in 22 publications with 33 experiments for the contrast HC>MCI. However, only 16 publications (17 experiments) remained after a more careful inspection (Table S2, Fig.1). For the contrast HC>AD, the initial search resulted in 71 publications, with 103 experiments but after a detailed inspection only 37 publications (37 experiments) could be included in the analysis (Table S2, Fig. 1).

**Figure 1.**
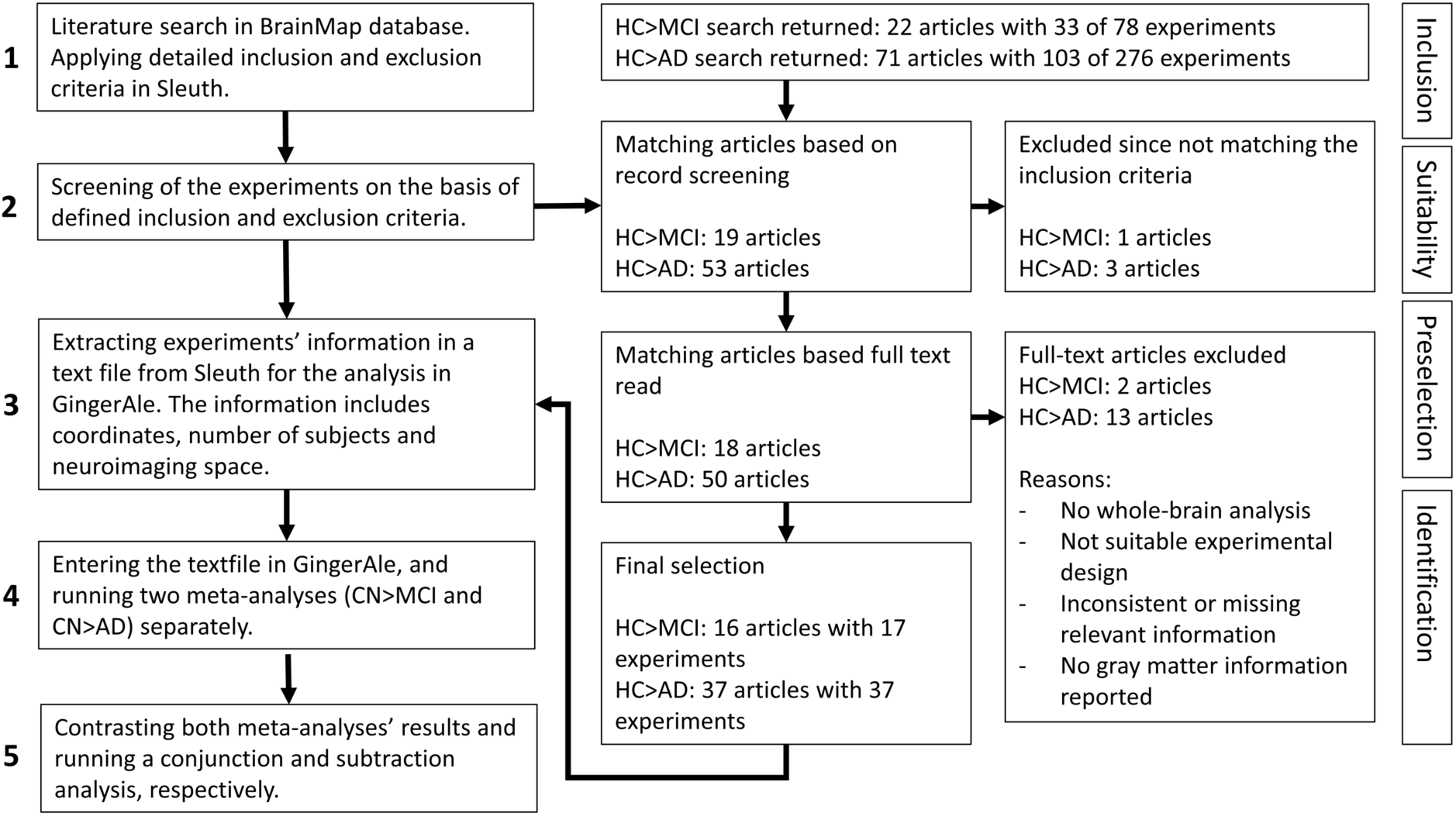
Flowchart of the steps performed in the meta-analysis.

**Table 1:** Inclusion and exclusion criteria for the systematic literature research.

| Inclusion criteria | Exclusion criteria |
| --- | --- |
| Study contrasted MCI vs HC or AD vs HC. | Study did not contrast MCI vs HC or AD vs HC, e.g., MCI vs AD. |
| MCI/AD and HC were matched regarding age, gender, education etc. | MCI/AD and HC were not matched regarding age, gender, education etc. or compared with patients other than MCI/AD. |
| Patient groups were clearly defined and if necessary summarized, e.g., “early AD” or “late AD”. | Several subgroups of AD were defined (e.g., early/late AD) and analyzed separately. |
| Study was based on one temporal measurement per patient. In longitudinal studies, the first MRI acquisition was selected. | Multiple measurements across time for each subject included in one analysis. |
| Analysis and report of gray matter based whole brain VBM. | Other analysis than gray matter based whole brain VBM, e.g., region of interest (ROI) analysis or correlation analysis. |

In total, structural MRI data from 347 MCI patients with 462 matching controls, and 799 AD patients with 973 matching controls were analyzed. Therefore, data from 1146 patients were compared with 1435 healthy older adults. A possible sample overlap between the studies was avoided by applying appropriate exclusion criteria in BrainMap, and additional cross-checks with the primary literature. Specifically, this included the sample characteristics, author affiliations and scanning locations. Note, however, that in the HC>MCI meta-analysis, two experiments from Bozzali et al. (2006) were included, contrasting MCI retrospective converters and non-converters with healthy controls separately. Here, these experiments were treated as independent analyses due to different MCI groups involved, the focus on coordinate-based contrasts, and no other coding possibility of the experiments (Turkeltaub et al., 2012). Additionally, in four studies (Bozzali et al., 2006; Hämäläinen et al., 2007; Rami et al., 2009; Shiino et al., 2006) the same HCs were contrasted against MCI and AD, respectively; here they were entered into both meta-analyses HC>MCI and HC>AD, respectively. However, study-specific biases are unlikely due to the number of included experiments (see Figure 1) (Heckner et al., 2021; Turkeltaub et al., 2012).

MRI scanners included field strengths from 1.0 to 3.0 Tesla, for one study the field strength was unknown, and all incorporated studies performed a VBM whole brain analysis (Ashburner and Friston, 2000; Mechelli et al., 2005), typically with SPM, MedX or FSL. A possible bias resulting from different MRI scanner field strengths (Tardif et al., 2009) was investigated using a chi-squared test. It revealed no significant effects, indicating no systematic differences in MRI field strength in the two meta-analyses (χ²(3) = 1.55, p = 0.67, Cramer’s V= 0.168).

About 80 percent of the studies reported their findings in MNI and 20 percent in Talairach reference space. Ten of 17 experiments used voxel-or cluster-wise corrected p-values in the HC>MCI comparison, the remaining seven used uncorrected p-values. In the HC>AD comparison 30 of 37 experiments used corrected p-values and seven used uncorrected p-values. All information is listed in detail in Table S2 and Table S3.

Two researchers (M.M. and H.M.) independently inspected each study individually with regard to the inclusion and exclusion criteria, based on the abstract, title, information provided by Sleuth, and finally, a full text read. Disagreements were, if required, discussed with a third researcher (N.B.). The final selection of included publications and experiments, respectively, with further study-specific information is shown in Tables S2 and S3.

### Statistical analysis

#### Activation likelihood estimation (ALE)

The activation likelihood estimation (ALE) meta-analytic approach was used to investigate the convergence of VBM results from all included whole-brain experiments (Eickhoff et al., 2012, 2009; Turkeltaub et al., 2012; Vanasse et al., 2018). ALE is implemented in BrainMap’s application GingerALE (version 3.0.2) (Eickhoff et al., 2012, 2009; Turkeltaub et al., 2012) and represents a coordinate-based neuroimaging meta-analyses technique. Here, in a first step, all reported Talairach coordinates were automatically converted to MNI space by the implemented algorithm icbm2tal (Laird et al., 2010; Lancaster et al., 2007). All other studies, i.e., those that already provided MNI coordinates, were not further converted. Second, the ALE random-effects algorithm treated all included coordinates of maximum activation, called foci, as spatial Gaussian probability distributions weighted by the number of subjects (Eickhoff et al., 2009; Laird et al., 2009). Further, the probability distributions of the foci from the specific experiment are summarized per voxel, resulting in a model activation (MA) map. Here, the voxel-wise MA values were calculated based on the maximum probability of any focus in the given experiment (Eickhoff et al., 2012; Turkeltaub et al., 2012). The voxel-wise union of these MA maps results in ALE scores on a voxel level. Accordingly, ALE scores represent the convergence of results across studies instead of foci. Third, to statistically evaluate the degree of convergence across studies, ALE scores are compared to an empirical null distribution of random spatial associations between all MA maps (Eickhoff et al., 2012, 2009). Here, our results were thresholded at p<0.05 with 1,000 permutations (cluster-level) corrected for multiple comparisons by the family-wise error (FWE) and an cluster-forming (uncorrected) p<0.001 (Eickhoff et al., 2016). Again, this approach was conducted separately for the meta-analysis HC>MCI and HC>AD.

Additionally, to investigate the differences and common effects of the two meta-analyses HC>MCI and HC>AD, their results were combined in a contrast analysis ([HC>MCI]>[HC>AD] and [HC>AD]>[HC>MCI]) and a conjunction analysis (HC>MCI and HC>AD), respectively. Therefore, the conjunction included computing the voxel-wise-minimum based on the CN>MCI and CN>AD ALE images resulting from initial meta-analysis (Eickhoff et al., 2011; Laird et al., 2005). The subtraction resulted in a new ALE image for each dataset, which was then subtracted from the other and compared to the true data. Finally, after permutations, a final voxel-wise p-value image is created and then z-scored to represent significance rather than the direct ALE subtraction (Eickhoff et al., 2011; Laird et al., 2005). Note that different sample sizes between the meta-analyses were corrected for the conjunction and contrast (Eickhoff et al., 2011). The results were thresholded at p<0.01 (uncorrected) and 10,000 permutations.

Significant clusters and associated brain regions will be reported based on the probabilistic cytoarchitectonic human brain maps as implemented in the SPM Anatomy Toolbox Version 3.0 (Eickhoff et al., 2007, 2006, 2005; Heckner et al., 2021) (available from https://www.fz-juelich.de/en/inm/inm-7/resources/jubrain-anatomy-toolbox).

#### Region-of-Interest analyses

On the basis of our a priori hypotheses (see introduction), we performed post-hoc region-of-interest (ROI) analyses for the NbM and EC, separately, using the whole-brain ALE meta-analysis approach. The ROIs for both regions were generated for the left and right hemisphere as well as one bilateral ROI for the NbM and one bilateral ROI for the EC. This was done in MNI space using the SPM Anatomy Toolbox Version 3.0 (Eickhoff et al., 2007, 2006, 2005) (available from https://www.fz-juelich.de/en/inm/inm-7/resources/jubrain-anatomy-toolbox). Specifically, the NbM was defined using previous published probabilistic maps labeled as Ch4 (Zaborszky et al., 2008), and the EC was derived from the previously published probability map (Amunts et al., 2005). The mean p-values of the NbM and EC were extracted at a threshold of 50% probability from the uncorrected p-value maps (resulting from the analyses HC>MCI and HC>AD, respectively) using Marsbar (Brett et al., 2002) implemented in the SPM toolbox (SPM 12, https://www.fil.ion.ucl.ac.uk/spm/) for MATLAB®. Note that this approach allows the extraction of p-values from each ROI but no other statistical values such as standard deviations or individual data points.

## Results

The HC>MCI meta-analysis included 17 experiments from 16 studies, and the HC>AD meta-analysis included 37 experiments from 37 studies. The mean age of the subjects ranged from 62 to 76 years (HC>MCI) and 59 to 83 years (HC>AD), respectively. For both meta-analyses, the patient groups were matched to the HC regarding age and gender. See Table S2 and S3, respectively, for further information on sociodemographics.

### HC > MCI

The meta-analysis HC>MCI revealed two significant clusters (Table 2, Figure 2A). The first cluster was driven by seven studies and included the right amygdala and right hippocampus. The second cluster was driven by four studies and included the left hippocampus and left amygdala.

**Figure 2.**
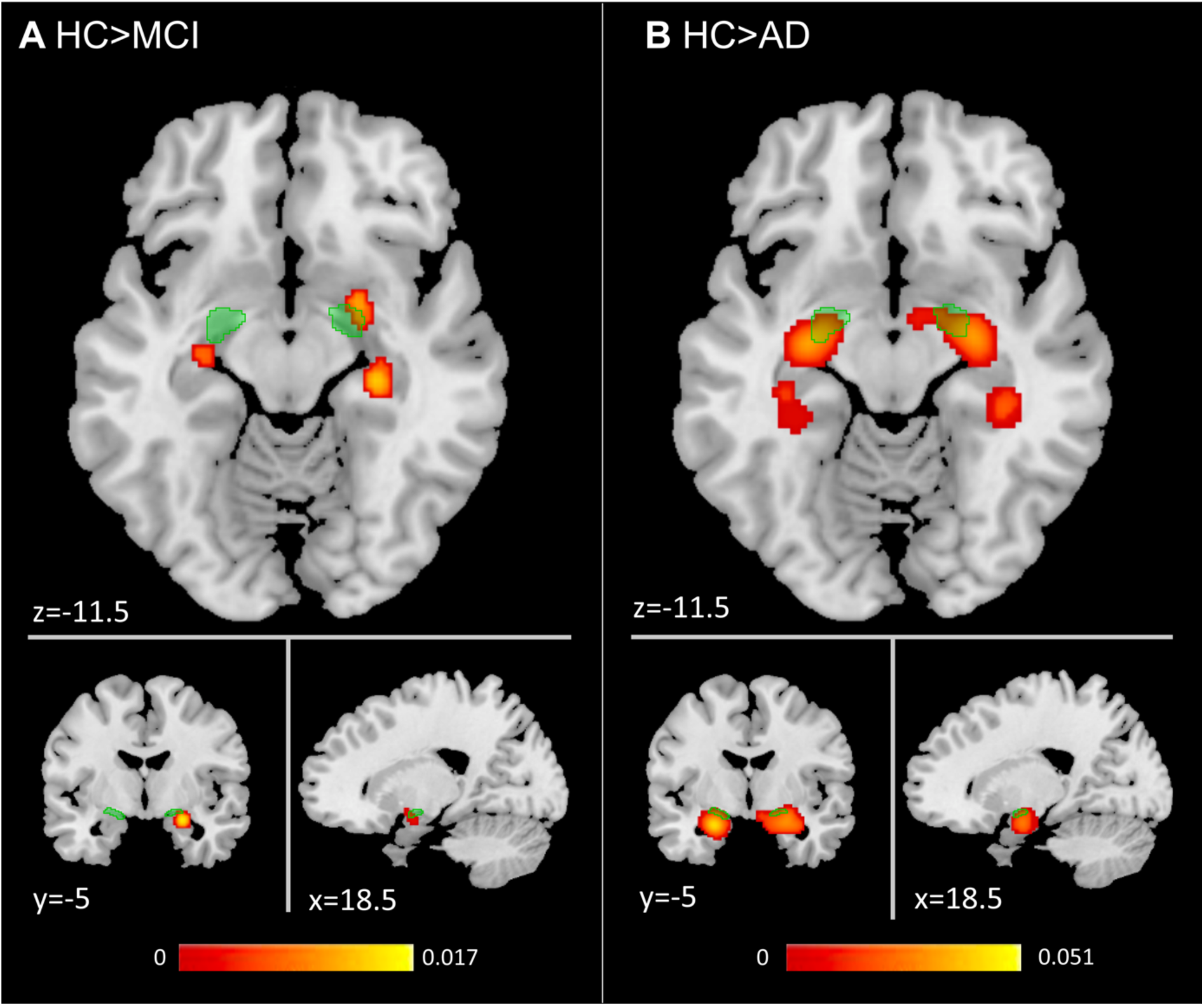
Results of meta-analyses HC>MCI (A) and HC>AD (B). Clusters represent significant convergence across experiments in gray matter reductions. Cluster-level p<0.05, family-wise error corrected with a cluster-forming threshold p<0.001. The color bar represents the activation likelihood estimation (ALE) scores. The light green region indicates the NbM. Clusters were overlaid onto a T1 weighted template image.

**Table 2:**
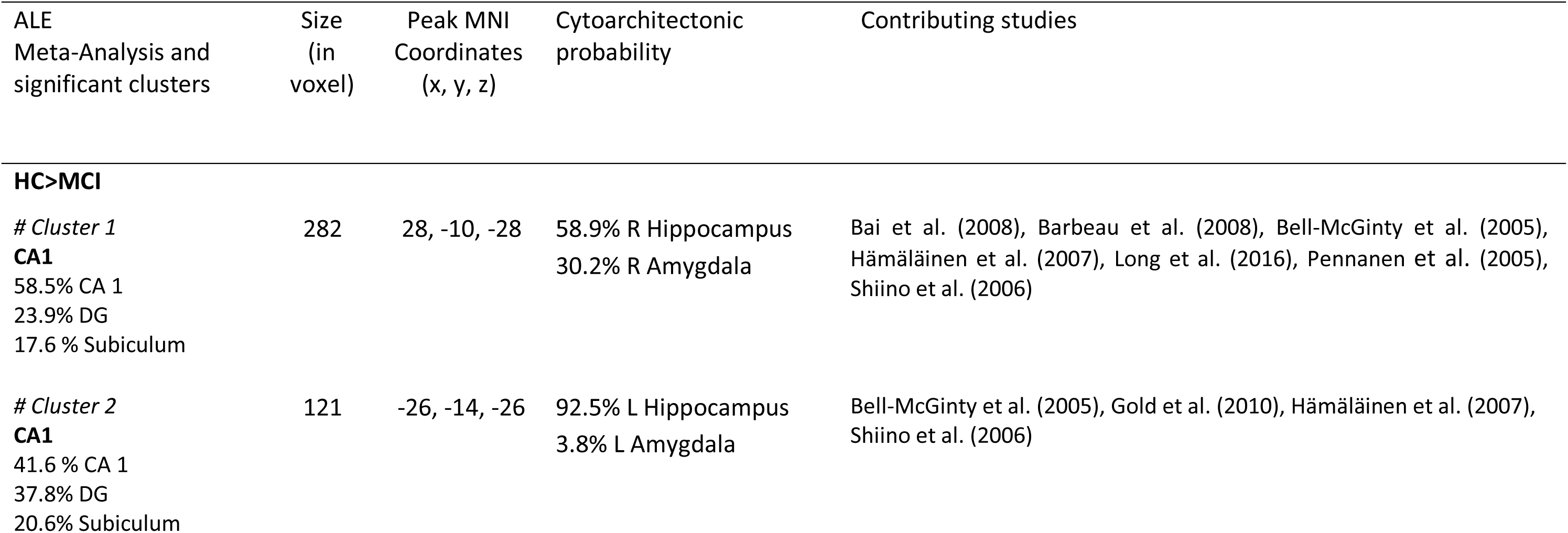
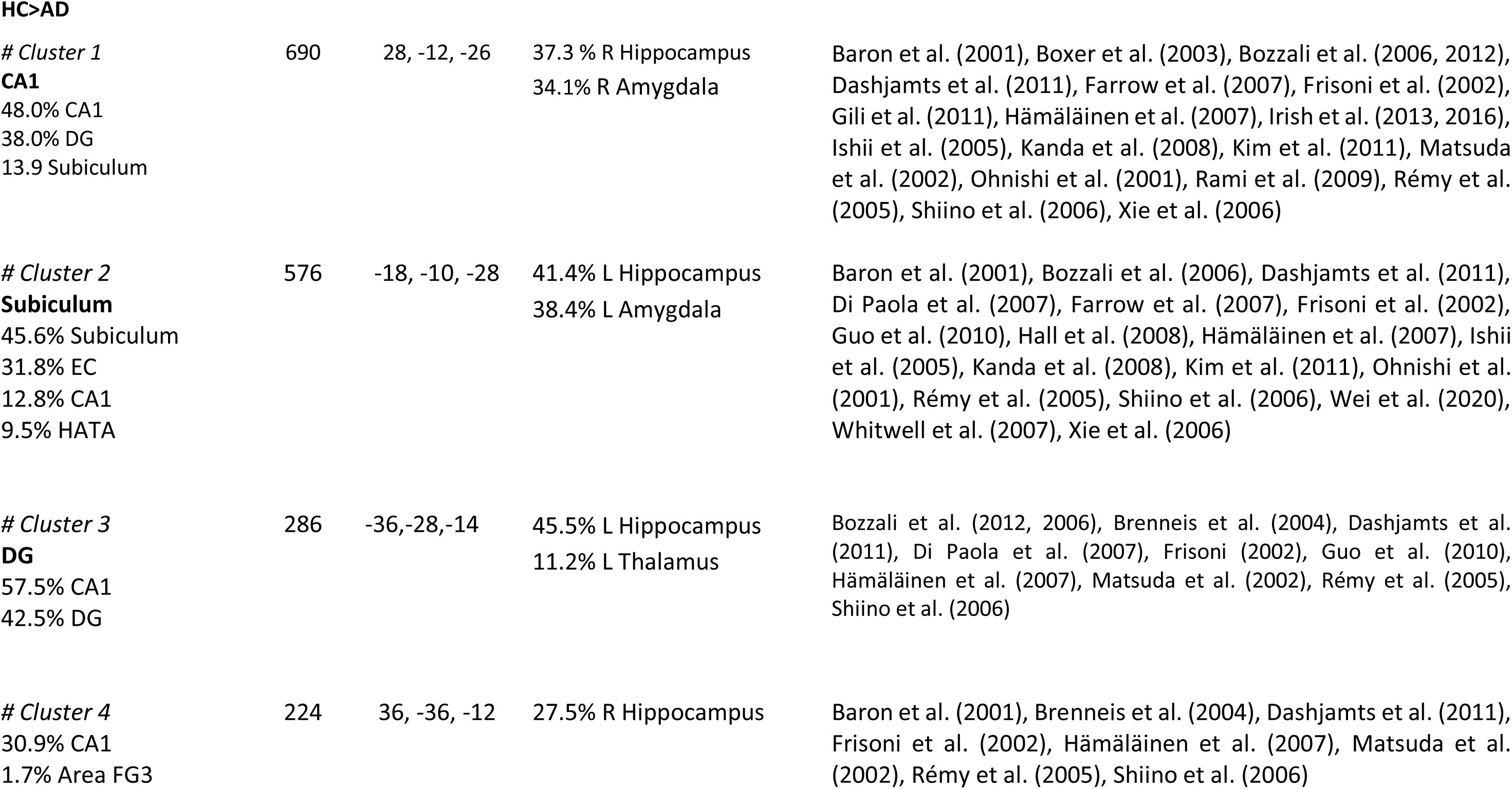
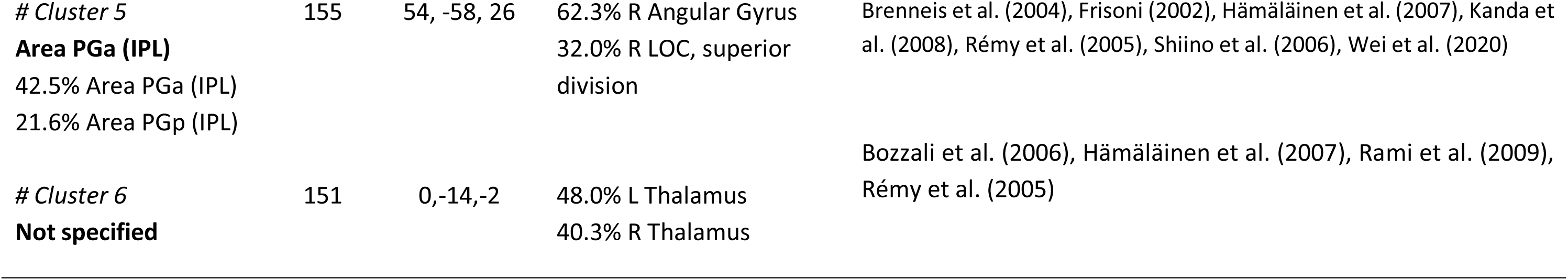
Results of the ALE meta-analysis for the contrasts HC>MCI and HC>AD. Significant clusters, their sizes (in voxels, 2×2×2 mm), peaks, regional assignment (probability in %) and cytoarchitectonic probabilities (in %) are based on the Jülich Brain atlas. Additionally, we also report the contributing studies, based on GingerALE. Abbreviations, R: Right; L: Left; CA: Cornu Ammonis; DG: Dentate Gyrus; HATA: Hippocampus-Amygdala-Transition Area; EC: Entorhinal Cortex; LOC: Lateral Occipital Cortex; Area FG3=Fusiform Gyrus; Area PGa (IPL)= Rostral region in the Angular Gyrus (Inferior Parietal Lobe); Area PGp (IPL)= Caudal region in the Angular Gyrus (Inferior Parietal Lobe)

### HC > AD

The meta-analysis HC>AD revealed six significant clusters (Table 2, Figure 2B). The first cluster was driven by twenty studies and included the right amygdala and right hippocampus. The second cluster was driven by eighteen studies and included the left amygdala, left hippocampus and EC. The third cluster was driven by eleven studies and included the left hippocampus and left thalamus. The fourth cluster was driven by eight studies and included the right hippocampus. The fifth cluster was driven by seven studies and included the right angular gyrus and right lateral occipital cortex (LOC). The sixth cluster was driven by four studies and included the left and right thalamus.

### ROI analysis

To further investigate possible gray matter differences in the NbM, a more liberal post-hoc ROI analysis based on the uncorrected p-value maps was carried out. It revealed a significant effect for the comparison HC>AD (p=0.007, Table 4) but not HC>MCI (p=0.122, Table 4). For the sake of completeness, in Table 4, we also report the p-values for the left and right NbM, which indicates a borderline significant effect in the right but not left NbM (p=0.073) for the comparison HC>MCI. In the EC, the analyses revealed no significant effects for HC>MCI (p=0.223) and HC>AD (p=0.208).

### Contrast and conjunction analyses

The conjunction of both meta-analyses (HC>AD & HC>MCI) showed two significant clusters. The first was driven by eleven experiments and included the right amygdala and right hippocampus (Table 3, Figure 3A). The second cluster was driven by nine experiments and included in the left hippocampus and left amygdala (Table 3, Figure 3A).

**Figure 3.**
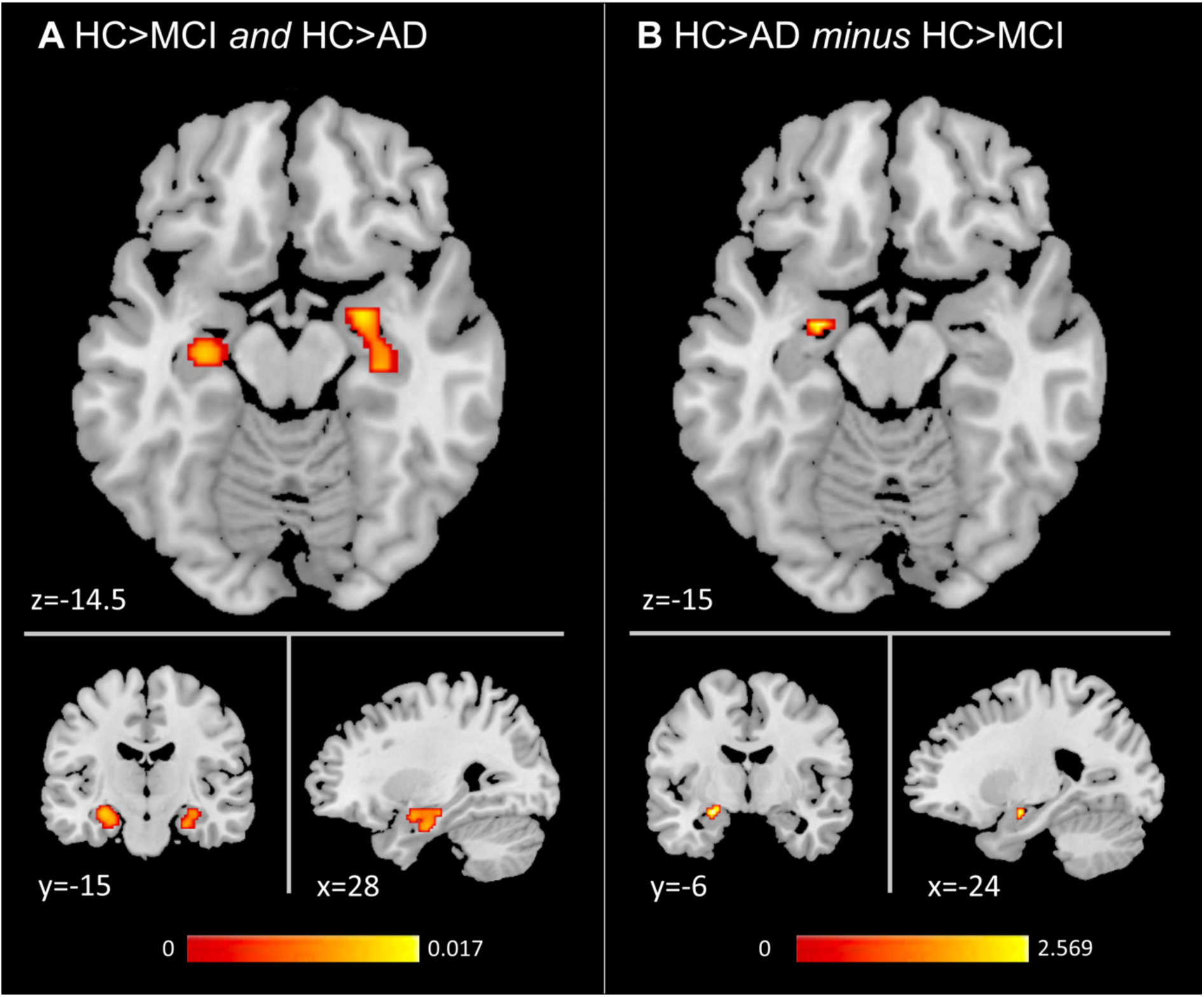
Results of the conjunction (A) and contrast analyses (B). A) shows the conjunction image (p<0.01, uncorrected) of both meta-analyses (HC>MCI and HC>AD), and color bar indicating the activation likelihood estimation scores. B) shows the results of the subtraction HC>AD *minus* HC>MCI (p<0.01, uncorrected), and the color bar reflects the z scores. Clusters were overlaid onto a T1 weighted template image.

**Table 3:**
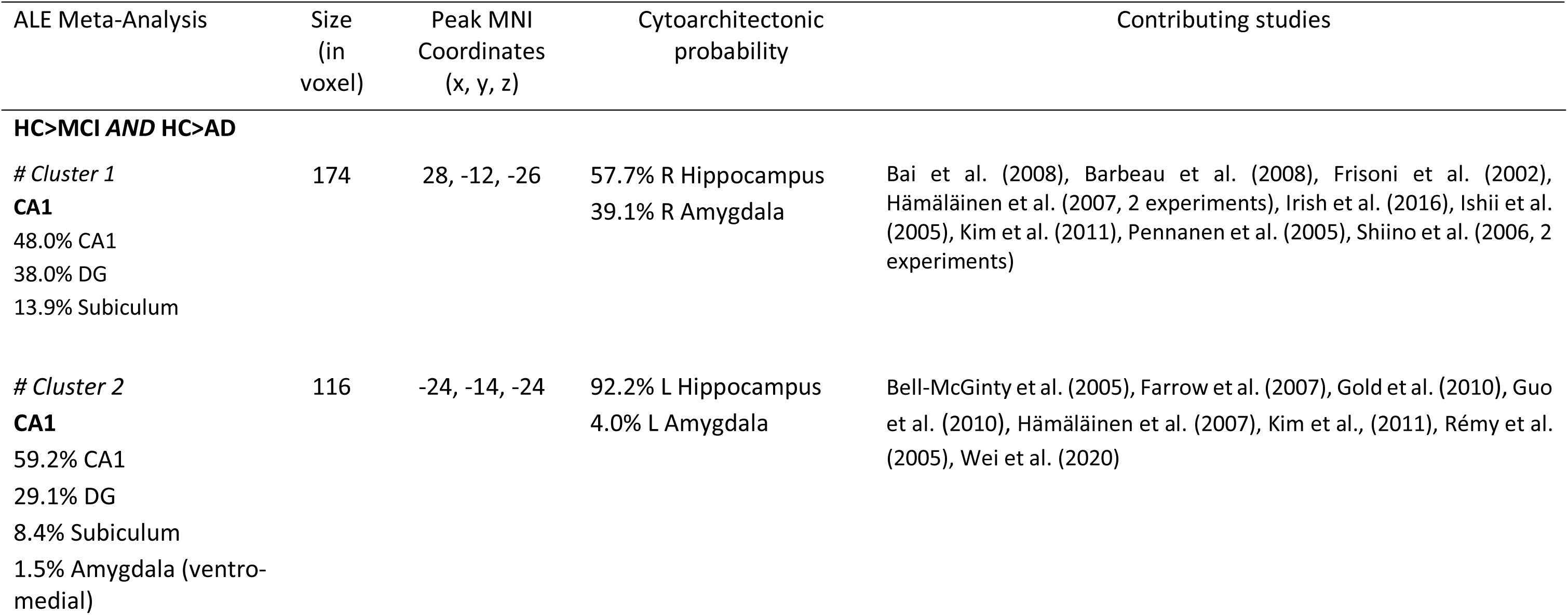
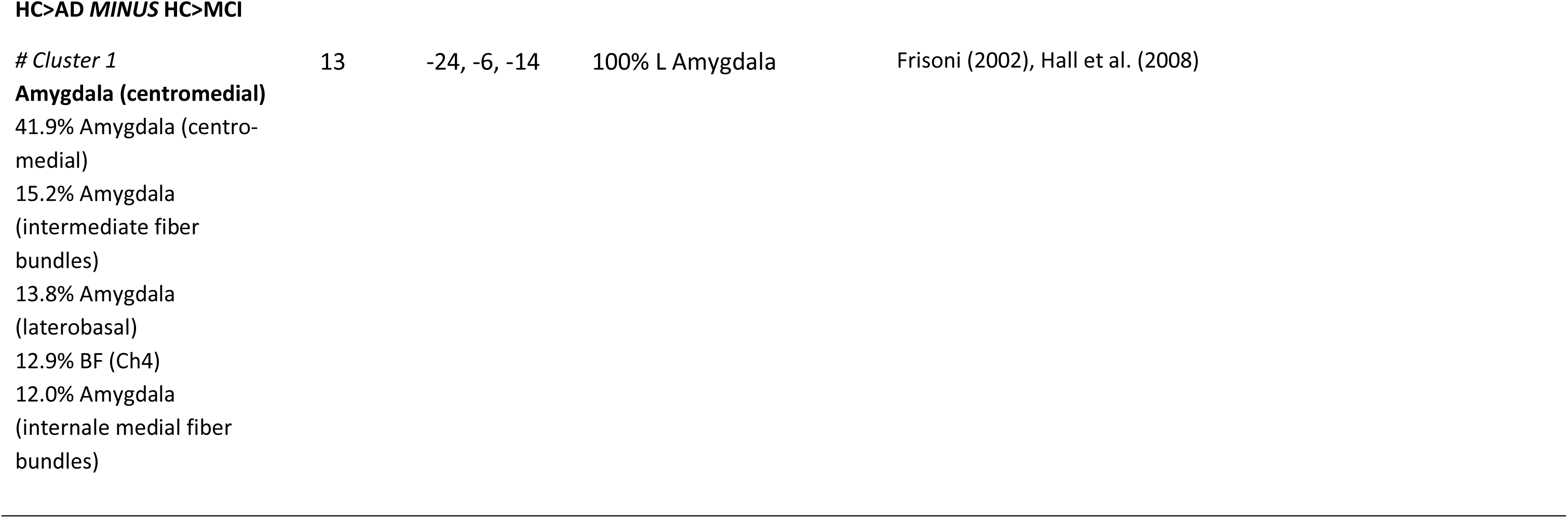
Results of the meta-analyses for the conjunction HC>MCI *AND* HC>AD as well as the contrast HC>MCI *MINUS* HC>AD. Significant clusters, their sizes (in voxels, 2×2×2 mm), peaks, regional assignment (probability in %) and cytoarchitectonic probabilities (in %) are based on the Jülich Brain atlas. Additionally, we also report the contributing studies, based on GingerALE. Abbreviations, R: Right; L: Left; CA: Cornu Ammonis; DG: Dentate Gyrus; BF: Basal Forebrain

Finally, the contrast of both meta-analyses (HC>AD minus HC>MCI) revealed a significant effect in the left amygdala (Table 3, Fig. 3B), which was driven by two studies. This indicates more pronounced gray matter differences in AD (<HC) as compared to MCI (<HC). The reverse contrast (HC>MCI minus HC>AD) revealed no significant effects.

## Discussion

We investigated structural abnormalities, with a focus on the cholinergic BF (NbM) and MTL, in MCI and AD patients compared to HC using the whole brain ALE meta-analytic approach. Our analyses included 53 previously published whole brain MRI studies with 54 experiments, and revealed less gray matter in the NbM of AD patients but only weak evidence in MCI. Furthermore, MCI and AD both showed, compared to HC, reduced gray matter in the MTL, including the hippocampus and amygdala, with hints towards more pronounced amygdala effects in AD. As such, our findings provide novel evidence in favor of the notion that structural degeneration of the cholinergic BF and interconnected MTL play a critical role in the progression of AD.

The cholinergic BF is a core region involved in complex behaviors and cognition, including attention and learning, through the modulation of neural activity in distant brain regions (Picciotto et al., 2012; Záborszky et al., 2018). It is also known to degenerate during healthy aging and AD, as shown in post mortem and *in vivo* imaging studies. For instance, in healthy older adults, the BF integrity, as measured with MRI based magnetization transfer ratio (MTR), closely related to verbal learning and working memory performance (Düzel et al., 2010). Importantly, in AD these structural changes appear to be much more pronounced, as reported in individual MRI studies (Hall et al., 2008; Whitwell et al., 2007), and evident in our post hoc ROI analysis based on the whole-brain estimated uncorrected p-value maps (Table 4), which further points towards a critical role of the BF, especially the NbM, in the disease progression. Indeed, post-mortem studies in AD have identified pathological changes of the NbM, including accumulations of amyloid-β and pTau, as early features that are associated with cognitive decline (Contestabile, 2011; Mesulam, 2004; Schliebs and Arendt, 2011). This selective vulnerability might be explained by the large projecting axons with wide arbors expanding vast distances in the central nervous system (Li et al., 2018; Mesulam and Geula, 1988; Wu et al., 2014). With regard to AD progression, previous studies suggest that pathologies of the BF precede those within other cortical regions, especially the EC and hippocampus. In humans, such a view is supported by longitudinal MRI studies showing that the BF volume predicts atrophy in the EC, which was further moderated by AD specific proteinopathies (Fernández-Cabello et al., 2020; Schmitz and Spreng, 2016). Finally, there is also evidence for reduced NbM volumes in patients with MCI and preclinical AD, which underlines the notion of a characteristic disease progression (Grothe et al., 2012; Hall et al., 2008; Kilimann et al., 2014; Muth et al., 2010).

**Table 4:** Results of the post-hoc ROI analysis. Listed are the average p-values extracted from the uncorrected p-value map of both meta-analyses (HC>MCI and HC>AD) for both the Nucleus basalis of Meynert (NbM) and Entorhinal cortex (EC). * indicates p<0.01.

| Contrast and ROI | p-value |
| --- | --- |
| **HC>MCI** |  |
| L NbM | 0.165 |
| R NbM | 0.073 |
| **Bilateral NbM** | 0.122 |
| L EC | 0.335 |
| R EC | 0.089 |
| **Bilateral EC** | 0.223 |
| **HC>AD** |  |
| L NbM | 0.012 |
| R NbM | 0.001* |
| **Bilateral NbM** | 0.007* |
| L EC | 0.121 |
| R EC | 0.312 |
| **Bilateral EC** | 0.208 |

In our meta-analysis, NbM gray matter reductions in MCI were only marginally significant (p=0.073, Table 4), which is contrary to our hypothesis and the pathological staging model (Fernández-Cabello et al., 2020; Schmitz and Spreng, 2016). While this is difficult to interpret, at least two explanations seem to be relevant. First, the number of included experiments for the meta-analysis HC>MCI (n=17) was much smaller as compared to HC>AD (n=37). Therefore, the null-finding could simply reflect a power problem, which could be addressed in future studies by searching more or other databases. Second, since NbM atrophy in MCI is not as pronounced as in AD, it is much more difficult to detect it with MRI based VBM. Indeed, not the entire NbM but posterior parts of it seem to be particularly vulnerable to the early stages of AD (Arendt et al., 1985; Doucette et al., 1986; Grothe et al., 2013, 2012; Kilimann et al., 2014; Liu et al., 2015; Vogels et al., 1990), which further calls for more fine-grained microstructural MRI measures such as multiparameter mapping (Leutritz et al., 2020) (see below).

Compared to healthy controls, both patient groups showed significant gray matter reductions in the amygdala with cautious evidence for more pronounced effects in AD (Table 3, Figure 3). This is, again, in line with individual studies that drive the effect (Hall et al., 2008; Kim et al., 2011) and others that also show early structural degenerations within the amygdala (Planche et al., 2022; Poulin et al., 2011; Ramos Bernardes da Silva Filho et al., 2017). From an anatomical point of view, the amygdala represents a central target region of NbM projections, including a high density of cholinergic axons (Mesulam and Geula, 1988; Mesulam et al., 1992), and functionally it is prominent for its role in processing emotional information (Dolan, 2002; LeDoux, 2003) but also memory formation and consolidation (Rutishauser et al., 2010; Yousuf et al., 2021). For instance, cholinergic stimulation in healthy middle aged humans reduced activity in the right amygdala for successful versus unsuccessful spatial context retrieval (Kukolja et al., 2009), which is in line with others suggesting that high levels of ACh promote encoding but interfere with retrieval (Winters et al., 2006). With regard to AD, a longitudinal study in humans could show that amygdala degeneration and abnormal memory-related alpha oscillations, as measured with electroencephalography (EEG), predict the conversion to AD (Prieto del Val et al., 2016). Further, fMRI based functional connectivity between the NbM and amygdala during resting state was increased with neuropathology as indicated by amyloid and tau PET (Zeng et al., 2022). This is in line with other PET studies on the spread of amyloid-β and tau (Insel et al., 2020), as well as MRI studies in MCI and AD showing that NbM atrophy covaries with atrophy in innervating regions, including the amygdala (Cantero et al., 2016; Schmitz et al., 2018). Therefore, the loss of cholinergic inputs to the amygdala, together with a trans-synaptic spread of tau, might facilitate and accelerate AD pathology (Clavaguera et al., 2009; de Calignon et al., 2012; Khan et al., 2014).

As expected, both meta-analyses (HC>MCI and HC>AD) revealed significant hippocampal gray matter atrophy in AD and MCI compared to HC. However, in the EC a significant effect was observed only in AD (Table 2, second cluster, 31.8% probability based on the Jülich Brain atlas) and not MCI. Note that our post hoc ROI analysis did not confirm the effect in AD possibly since it was spatially limited to a small portion of the rather large EC. In any case, these findings are only partly compatible with another prevailing model suggesting the trans-entorhinal cortex as origin of AD pathology (Braak and Braak, 1991; Corder et al., 2000; Duyckaerts et al., 1990). Together with an arguably stronger effect in the NbM in AD, this could be interpreted as evidence in favor of the view that NbM pathology precedes EC pathology (e.g., Schmitz and Spreng, 2016; Fernández-Cabello et al., 2020). However, the absence of structural degenerations in EC might also have methodological reasons, see below. Therefore, more fine-grained MRI based methods should be used, preferably in combination with larger samples and longitudinal designs, to further address this important issue.

The significant gray matter effect in the hippocampus in AD and MCI was driven by large numbers of studies (Table 3) and confirms, on a meta-analytic level, its involvement in AD pathology and its important role in early AD detection (Good et al., 2002; Hett et al., 2019; Planche et al., 2022; Wolz et al., 2011). With regard to the BF, the hippocampus (and EC) receives a high density of cholinergic axons, which mainly originate in the medial septum and vertical diagonal band nuclei (i.e. Ch1 and Ch2, Mesulam and Geula, 1988; Mesulam et al., 1992); this is in contrast to the amygdala, which receives cholinergic innervations from NbM (i.e. CH4) (Mesulam et al., 1983). Here, we focused in the NbM due to its early vulnerability in AD but the hodological separation between BF nuclei and specific projection sites in the MTL points towards another fruitful avenue in future research to fully understand healthy and pathological age-related development.

Apart from cortical projections, the cholinergic system is interconnected with subcortical regions that are part of the dopaminergic mesolimbic system and play a central role in the encoding and consolidation of novel information into long-term memory (Bunzeck et al., 2014; Lisman et al., 2011; Lisman and Grace, 2005). For instance, substantia nigra / ventral tegmental area (SN/VTA) dopamine neurons receive cholinergic projections from the laterodorsal and pedunculopontine (PPT) tegmental nuclei, which modulate their activity via muscarinic and nicotinic receptors (Futami et al., 1995; Oakman et al., 1995). Therefore, systemic injections of the nonselective muscarinic antagonist scopolamine increased dopamine release into the striatum in rats (Chapman et al., 1997; Miller and Blaha, 2004), and cholinergic agonists, and the other hand, attenuated the striatal dopamine efflux (Miller and Blaha, 2004). In humans, both the dopamine precursor levodopa and the acetylcholinesterase inhibitor galantamine modulated hemodynamic activity in the SN/VTA and MTL during novelty processing (Bunzeck et al., 2014). Therefore, these and other studies (e.g. Mark et al., 2011; Threlfell et al., 2012) point towards a close structural and functional interaction between the cholinergic and dopaminergic system.

Importantly and similar to the cholinergic system, the dopaminergic mesolimbic system also undergoes age-related changes (Backman et al., 2006; Guitart-Masip et al., 2016) but their common effect on cognition and behavior across the life span remain little understood. For instance, the structural integrity of the SN/VTA, as measured with MRI based magnetization transfer ration, was reduced in healthy older adults (Bunzeck et al., 2007) and correlated with learning abilities (Düzel et al., 2008). Moreover, age-related iron accumulation and demyelination in basal ganglia structures are closely related to verbal memory (Steiger and Bunzeck, 2017) and executive functioning (Biel et al., 2021). Along these lines, microstructural brain changes, in particular iron depositions, are not only a hallmark of healthy but also pathological aging. In fact, excessive iron depositions have long been observed in AD and Parkinson’s Disease (PD) (Guan et al., 2022; Loeffler et al., 1995; Sian-Hülsmann et al., 2011), which points towards a dysfunction of brain iron homeostasis (Ward et al., 2014). From a mechanistic point of view, this could have neurotoxic effects via oxidative stress, which accelerates neurodegeneration (Ward et al., 2014). More specifically, enhanced iron levels have been linked to amyloid-! and tau in AD (Rogers et al., 2008; Yamamoto et al., 2002), and “-synuclein and Lewy body aggregation in PD (Ostrerova-Golts et al., 2000).

Together, cognitive deficits during healthy and pathological aging can have complex etiologies involving both the cholinergic and dopaminergic system (Amalric et al., 2021). Therefore, future studies in humans might leverage recent advances in microstructural MRI and PET imaging to focus on several open questions, including interactions. Another line of research might focus on functional brain changes in AD, which often precede structural degenerations (Johnson et al., 2012; Sperling, 2011; Warren and Moustafa, 2023) and could involve compensational mechanisms (Cabeza et al., 2018) to promote cognition.

Finally, our meta-analytic approach has several strengths but also limitations, which both could guide future research. First, we focused on the convergence across experiments on the basis of previously published whole-brain VBM studies that are included in the BrainMap database. Currently, these are limited to ca 20-30% of all published neuroimaging studies. While this may be considered a limitation, the advantage is that BrainMap’s database allows a fast and less error-prone search and data extraction (Müller et al., 2018). As such, it represents a powerful tool to provide robust and generalizable results (Heckner et al., 2021). The disadvantage of our whole brain approach is that we could not integrate original studies with a region of interest approach showing significant structural changes in the BF (e.g., Arrondo et al., 2022; Colloby et al., 2014; Fernández-Cabello et al., 2020; Jessen et al., 2006; Kerbler et al., 2015; Kilimann et al., 2014; Schmitz et al., 2018; Schmitz and Spreng, 2016) or reduced structural integrity of cholinergic white matter pathways in AD (Schumacher et al. 2022, Nemy et al., 2023).

Second, the NbM is a rather small brain structure but not all original studies were specifically optimized with regard to spatial resolution and imaging analysis. For instance, too large voxels in combination with inappropriate smoothing kernels might „wash-out“ possible effects. Nevertheless, it is important to note that the voxel size (typically 1×1×1 mm or slightly larger) and smoothing kernel (approximately 8-12 mm full width at half maximum) employed in most studies should be sufficient to detect signals in the NbM. Third, the meta-analysis CN>MCI only included 17 experiments, which is at the lower end of recommendations (17-20) to achieve adequate power to detect even small effects (Eickhoff et al., 2016). Fourth, the contrast and conjunction analyses did not include a minimum cluster size (Eickhoff et al., 2012, 2011, 2009; Turkeltaub et al., 2012). This is particularly relevant for the rather small amygdala effect (Table 3, 13 voxels, which correspond to 104 mm^3^, driven by two studies), which, therefore, needs to be interpreted with caution.

Fifth, participants were included in the analyses solely based on their clinical status. To avoid possible heterogeneity (e.g. by including preclinical AD cases in the group of healthy controls), the AT(N) framework, which incorporates amyloid-beta (A), tau (T), and neurodegenerative (N), could have been used (Jack et al., 2018). However, in most studies, these markers were not assessed, and larger sample sizes would be necessary to include at least three groups (A-T-, A+T-, A+T+), with typically imbalanced distributions (see, e.g., Zeng et al. (2022)). Additionally, other forms of dementia were excluded based on the specified exclusion criteria in BrainMap that were validated by cross-checking the original literature. Finally, the direct comparison of MCI>AD (rather than HC>MCI vs HC>AD) might have been helpful, but a search in BrainMap’s database revealed only four studies, which is not suitable to perform a statistical meta-analysis (Eickhoff et al., 2016).

In conclusion, using the ALE meta-analytic approach, involving 54 experiments and 2581 subjects, our study provides novel evidence for structural degenerations of the NbM in AD patients, but only limited evidence in MCI. In both patient groups, the hippocampus and amygdala showed gray matter reductions, with hints towards more pronounced amygdala effects in AD. As such, our findings provide novel insights into the role of the cholinergic basal forebrain and interconnected MTL in AD.

## Acknowledgements

This research was financially supported by the University of Lübeck.

## References

Acquas, E., Wilson, C., Fibiger, H.C., 1996. Conditioned and Unconditioned Stimuli Increase Frontal Cortical and Hippocampal Acetylcholine Release: Effects of Novelty, Habituation, and Fear. J. Neurosci. 16, 3089–3096. 10.1523/JNEUROSCI.16-09-03089.1996

Aigner, T.G., Mishkin, M., 1986. The effects of physostigmine and scopolamine on recognition memory in monkeys. Behav Neural Biol 45, 81–7.

Amalric, M., Pattij, T., Sotiropoulos, I., Silva, J.M., Sousa, N., Ztaou, S., Chiamulera, C., Wahlberg, L.U., Emerich, D.F., Paolone, G., 2021. Where Dopaminergic and Cholinergic Systems Interact: A Gateway for Tuning Neurodegenerative Disorders. Front. Behav. Neurosci. 15.

Amunts, K., Kedo, O., Kindler, M., Pieperhoff, P., Mohlberg, H., Shah, N.J., Habel, U., Schneider, F., Zilles, K., 2005. Cytoarchitectonic mapping of the human amygdala, hippocampal region and entorhinal cortex: intersubject variability and probability maps. Anat. Embryol. (Berl.) 210, 343–352. 10.1007/s00429-005-0025-5

Arendt, T., Bigl, V., Tennstedt, A., Arendt, A., 1985. Neuronal loss in different parts of the nucleus basalis is related to neuritic plaque formation in cortical target areas in alzheimer’s disease. Neuroscience 14, 1–14. 10.1016/0306-4522(85)90160-5

Arendt, T., Brückner, M.K., Morawski, M., Jäger, C., Gertz, H.-J., 2015. Early neurone loss in Alzheimer’s disease: cortical or subcortical? Acta Neuropathol. Commun. 3, 10. 10.1186/s40478-015-0187-1

Arrondo, P., Elía-Zudaire, Ó., Martí-Andrés, G., Fernández-Seara, M.A., Riverol, M., 2022a. Grey matter changes on brain MRI in subjective cognitive decline: a systematic review. Alzheimers Res. Ther. 14, 98. 10.1186/s13195-022-01031-6

Arrondo, P., Elía-Zudaire, Ó., Martí-Andrés, G., Fernández-Seara, M.A., Riverol, M., 2022b. Grey matter changes on brain MRI in subjective cognitive decline: a systematic review. Alzheimers Res. Ther. 14, 98. 10.1186/s13195-022-01031-6

Ashburner, J., Friston, K.J., 2000. Voxel-based morphometry--the methods. Neuroimage 11, 805–21.

Atri, A., Sherman, S., Norman, K.A., Kirchhoff, B.A., Nicolas, M.M., Greicius, M.D., Cramer, S.C., Breiter, H.C., Hasselmo, M.E., Stern, C.E., 2004. Blockade of central cholinergic receptors impairs new learning and increases proactive interference in a word paired-associate memory task. Behav Neurosci 118, 223–36.

Backman, L., Nyberg, L., Lindenberger, U., Li, S.C., Farde, L., 2006. The correlative triad among aging, dopamine, and cognition: Current status and future prospects. Neurosci Biobehav Rev 30, 791–807.

Bai, F., Zhang, Z., Yu, H., Shi, Y., Yuan, Y., Zhu, W., Zhang, X., Qian, Y., 2008. Default-mode network activity distinguishes amnestic type mild cognitive impairment from healthy aging: A combined structural and resting-state functional MRI study. Neurosci. Lett. 438, 111–115. 10.1016/j.neulet.2008.04.021

Baker-Nigh, A., Vahedi, S., Davis, E.G., Weintraub, S., Bigio, E.H., Klein, W.L., Geula, C., 2015. Neuronal amyloid-β accumulation within cholinergic basal forebrain in ageing and Alzheimer’s disease. Brain 138, 1722–1737. 10.1093/brain/awv024

Ballinger, E.C., Ananth, M., Talmage, D.A., Role, L.W., 2016. Basal Forebrain Cholinergic Circuits and Signaling in Cognition and Cognitive Decline. Neuron 91, 1199–1218. 10.1016/j.neuron.2016.09.006

Barbeau, E.J., Ranjeva, J.P., Didic, M., Confort-Gouny, S., Felician, O., Soulier, E., Cozzone, P.J., Ceccaldi, M., Poncet, M., 2008. Profile of memory impairment and gray matter loss in amnestic mild cognitive impairment. Neuropsychologia 46, 1009–1019. 10.1016/j.neuropsychologia.2007.11.019

Baron, J.C., Chételat, G., Desgranges, B., Perchey, G., Landeau, B., de la Sayette, V., Eustache, F., 2001. In vivo mapping of gray matter loss with voxel-based morphometry in mild Alzheimer’s disease. NeuroImage 14, 298–309. 10.1006/nimg.2001.0848

Bauer, M., Kluge, C., Bach, D., Bradbury, D., Heinze, H.J., Dolan, R.J., Driver, J., 2012. Cholinergic Enhancement of Visual Attention and Neural Oscillations in the Human Brain. Curr. Biol. 22, 397–402. 10.1016/j.cub.2012.01.022

Bell-McGinty, S., Lopez, O.L., Meltzer, C.C., Scanlon, J.M., Whyte, E.M., Dekosky, S.T., Becker, J.T., 2005. Differential cortical atrophy in subgroups of mild cognitive impairment. Arch. Neurol. 62, 1393–1397. 10.1001/archneur.62.9.1393

Biel, D., Steiger, T.K., Bunzeck, N., 2021. Age-related iron accumulation and demyelination in the basal ganglia are closely related to verbal memory and executive functioning. Sci. Rep. 11, 9438. 10.1038/s41598-021-88840-1

Boxer, A.L., Rankin, K.P., Miller, B.L., Schuff, N., Weiner, M., Gorno-Tempini, M.-L., Rosen, H.J., 2003. Cinguloparietal atrophy distinguishes Alzheimer disease from semantic dementia. Arch. Neurol. 60, 949–956. 10.1001/archneur.60.7.949

Bozzali, M., Filippi, M., Magnani, G., Cercignani, M., Franceschi, M., Schiatti, E., Castiglioni, S., Mossini, R., Falautano, M., Scotti, G., Comi, G., Falini, A., 2006. The contribution of voxel-based morphometry in staging patients with mild cognitive impairment. Neurology 67, 453–460. 10.1212/01.wnl.0000228243.56665.c2

Bozzali, M., Giulietti, G., Basile, B., Serra, L., Spanò, B., Perri, R., Giubilei, F., Marra, C., Caltagirone, C., Cercignani, M., 2012. Damage to the cingulum contributes to Alzheimer’s disease pathophysiology by deafferentation mechanism. Hum. Brain Mapp. 33, 1295–1308. 10.1002/hbm.21287

Braak, H., Braak, E., 1991. Neuropathological stageing of Alzheimer-related changes. Acta Neuropathol. (Berl.) 82, 239–259. 10.1007/BF00308809

Brenneis, C., Wenning, G., Egger, K., Schocke, M., Trieb, T., Seppi, K., Marksteiner, J., Ransmayr, G., Benke, T., Poewe, W., 2004. Basal forebrain atrophy is a distinctive pattern in dementia with Lewy bodies. Neuroreport 15, 1711–4. 10.1097/01.wnr.0000136736.73895.03

Brett, M., Anton, J.L., Valabrgue, R., Poline, J.-B., 2002. Region of interest analysis using an SPM toolbox. Presented at the 8th International Conference on Functional Mapping of the Human Brain, June 2-6, 2002, Sendai, Japan. Neuroimage 13, 210–217.

Buccafusco, J.J., Letchworth, S.R., Bencherif, M., Lippiello, P.M., 2005. Long-lasting cognitive improvement with nicotinic receptor agonists: mechanisms of pharmacokinetic-pharmacodynamic discordance. Trends Pharmacol Sci 26, 352–60.

Bunzeck, N., Guitart-Masip, M., Dolan, R.J., Duzel, E., 2014. Pharmacological Dissociation of Novelty Responses in the Human Brain. Cereb. Cortex 24, 1351–1360. 10.1093/cercor/bhs420

Bunzeck, N., Schütze, H., Stallforth, S., Kaufmann, J., Düzel, S., Heinze, H.-J., Düzel, E., 2007. Mesolimbic Novelty Processing in Older Adults. Cereb. Cortex 17, 2940–2948. 10.1093/cercor/bhm020

Cabeza, R., Albert, M., Belleville, S., Craik, F.I.M., Duarte, A., Grady, C.L., Lindenberger, U., Nyberg, L., Park, D.C., Reuter-Lorenz, P.A., Rugg, M.D., Steffener, J., Rajah, M.N., 2018. Maintenance, reserve and compensation: the cognitive neuroscience of healthy ageing. Nat. Rev. Neurosci. 1. 10.1038/s41583-018-0068-2

Cantero, J.L., Zaborszky, L., Atienza, M., 2016. Volume Loss of the Nucleus Basalis of Meynert is Associated with Atrophy of Innervated Regions in Mild Cognitive Impairment. Cereb. Cortex cercor;bhw195v1. 10.1093/cercor/bhw195

Chapman, C.A., Yeomans, J.S., Blaha, C.D., Blackburn, J.R., 1997. Increased striatal dopamine efflux follows scopolamine administered systemically or to the tegmental pedunculopontine nucleus. Neuroscience 76, 177–86.

Clavaguera, F., Bolmont, T., Crowther, R.A., Abramowski, D., Frank, S., Probst, A., Fraser, G., Stalder, A.K., Beibel, M., Staufenbiel, M., Jucker, M., Goedert, M., Tolnay, M., 2009. Transmission and spreading of tauopathy in transgenic mouse brain. Nat. Cell Biol. 11, 909–913. 10.1038/ncb1901

Colloby, S.J., O’Brien, J.T., Taylor, J.-P., 2014. Patterns of cerebellar volume loss in dementia with Lewy bodies and Alzheimer’s disease: A VBM-DARTEL study. Psychiatry Res. 223, 187–191. 10.1016/j.pscychresns.2014.06.006

Contestabile, A., 2011. The history of the cholinergic hypothesis. Behav. Brain Res. 221, 334–340. 10.1016/j.bbr.2009.12.044

Corder, E.H., Woodbury, M.A., Volkmann, I., Madsen, D.K., Bogdanovic, N., Winblad, B., 2000. Density profiles of Alzheimer disease regional brain pathology for the Huddinge brain bank: pattern recognition emulates and expands upon Braak staging. Exp. Gerontol., Biological Aging - Euroconference on Molecular, Cellular and Tissue Gerontology 35, 851–864. 10.1016/S0531-5565(00)00147-9

Dashjamts, T., Yoshiura, T., Hiwatashi, A., Yamashita, K., Monji, A., Ohyagi, Y., Kamano, H., Kawashima, T., Kira, J.-I., Honda, H., 2011. Simultaneous arterial spin labeling cerebral blood flow and morphological assessments for detection of Alzheimer’s disease. Acad. Radiol. 18, 1492–1499. 10.1016/j.acra.2011.07.015

Davies, P., Maloney, A.J.F., 1976. Selective Loss of Central Cholinergic Neurons in Alzheimer’s Disease. The Lancet, Originally published as Volume 2, Issue 8000 308, 1403. 10.1016/S0140-6736(76)91936-X

de Calignon, A., Polydoro, M., Suárez-Calvet, M., William, C., Adamowicz, D.H., Kopeikina, K.J., Pitstick, R., Sahara, N., Ashe, K.H., Carlson, G.A., Spires-Jones, T.L., Hyman, B.T., 2012. Propagation of tau pathology in a model of early Alzheimer’s disease. Neuron 73, 685– 697. 10.1016/j.neuron.2011.11.033

de Lacalle, S., Cooper, J.D., Svendsen, C.N., Dunnett, S.B., Sofroniew, M.V., 1996. Reduced retrograde labelling with fluorescent tracer accompanies neuronal atrophy of basal forebrain cholinergic neurons in aged rats. Neuroscience 75, 19–27. 10.1016/0306-4522(96)00239-4

de Calignon, A., Polydoro, M., Suárez-Calvet, M., William, C., Adamowicz, D.H., Kopeikina, K.J., Pitstick, R., Sahara, N., Ashe, K.H., Carlson, G.A., Spires-Jones, T.L., Hyman, B.T., 2012. Propagation of Tau Pathology in a Model of Early Alzheimer’s Disease. Neuron 73, 685– 697. 10.1016/j.neuron.2011.11.033

Di Paola, M., Macaluso, E., Carlesimo, G.A., Tomaiuolo, F., Worsley, K.J., Fadda, L., Caltagirone, C., 2007. Episodic memory impairment in patients with Alzheimer’s disease is correlated with entorhinal cortex atrophy. A voxel-based morphometry study. J. Neurol. 254, 774–781. 10.1007/s00415-006-0435-1

Dolan, R.J., 2002. Emotion, Cognition, and Behavior. Science 298, 1191–1194. 10.1126/science.1076358

Doucette, R., Fisman, M., Hachinski, V.C., Mersky, H., 1986. Cell Loss from the Nucleus Basalis of Meynert in Alzheimer’s Disease. Can. J. Neurol. Sci. J. Can. Sci. Neurol. 13, 435–440. 10.1017/S0317167100037070

Duyckaerts, C., Delaère, P., Hauw, J.-J., Abbamondi-Pinto, A.L., Sorbi, S., Allen, I., Brion, J.P., Flament-Durand, J., Duchen, L., Kauss, J., Schlote, W., Lowe, J., Probst, A., Ravid, R., Swaab, D.F., Renkawek, K., Tomlinson, B., 1990. Rating of the lesions in senile dementia of the Alzheimer type: concordance between laboratories A European multicenter study under the auspices of EURAGE. J. Neurol. Sci. 97, 295–323. 10.1016/0022-510X(90)90226-D

Düzel, S., Münte, T.F., Lindenberger, U., Bunzeck, N., Schütze, H., Heinze, H.-J., Düzel, E., 2010. Basal forebrain integrity and cognitive memory profile in healthy aging. Brain Res. 1308, 124–136. 10.1016/j.brainres.2009.10.048

Düzel, S., Schütze, H., Stallforth, S., Kaufmann, J., Bodammer, N., Bunzeck, N., Münte, T.F., Lindenberger, U., Heinze, H.-J., Düzel, E., 2008. A close relationship between verbal memory and SN/VTA integrity in young and older adults. Neuropsychologia 46, 3042– 3052. 10.1016/j.neuropsychologia.2008.06.001

Eckart, C., Bunzeck, N., 2013. Dopamine modulates processing speed in the human mesolimbic system. NeuroImage 66, 293–300. 10.1016/j.neuroimage.2012.11.001

Eckart, C., Woźniak-Kwaśniewska, A., Herweg, N.A., Fuentemilla, L., Bunzeck, N., 2016. Acetylcholine modulates human working memory and subsequent familiarity based recognition via alpha oscillations. NeuroImage 137, 61–69. 10.1016/j.neuroimage.2016.05.049

Eickhoff, S.B., Bzdok, D., Laird, A.R., Kurth, F., Fox, P.T., 2012. Activation likelihood estimation meta-analysis revisited. NeuroImage 59, 2349–2361. 10.1016/j.neuroimage.2011.09.017

Eickhoff, S.B., Bzdok, D., Laird, A.R., Roski, C., Caspers, S., Zilles, K., Fox, P.T., 2011. Co-activation patterns distinguish cortical modules, their connectivity and functional differentiation. NeuroImage 57, 938–949. 10.1016/j.neuroimage.2011.05.021

Eickhoff, S.B., Heim, S., Zilles, K., Amunts, K., 2006. Testing anatomically specified hypotheses in functional imaging using cytoarchitectonic maps. NeuroImage 32, 570–582. 10.1016/j.neuroimage.2006.04.204

Eickhoff, S.B., Laird, A.R., Grefkes, C., Wang, L.E., Zilles, K., Fox, P.T., 2009. Coordinate-based activation likelihood estimation meta-analysis of neuroimaging data: A random-effects approach based on empirical estimates of spatial uncertainty. Hum. Brain Mapp. 30, 2907–2926. 10.1002/hbm.20718

Eickhoff, S.B., Nichols, T.E., Laird, A.R., Hoffstaedter, F., Amunts, K., Fox, P.T., Bzdok, D., Eickhoff, C.R., 2016. Behavior, sensitivity, and power of activation likelihood estimation characterized by massive empirical simulation. NeuroImage 137, 70–85. 10.1016/j.neuroimage.2016.04.072

Eickhoff, S.B., Paus, T., Caspers, S., Grosbras, M.-H., Evans, A.C., Zilles, K., Amunts, K., 2007. Assignment of functional activations to probabilistic cytoarchitectonic areas revisited. NeuroImage 36, 511–521. 10.1016/j.neuroimage.2007.03.060

Eickhoff, S.B., Stephan, K.E., Mohlberg, H., Grefkes, C., Fink, G.R., Amunts, K., Zilles, K., 2005. A new SPM toolbox for combining probabilistic cytoarchitectonic maps and functional imaging data. NeuroImage 25, 1325–1335. 10.1016/j.neuroimage.2004.12.034

Farrow, T.F.D., Thiyagesh, S.N., Wilkinson, I.D., Parks, R.W., Ingram, L., Woodruff, P.W.R., 2007. Fronto-temporal-lobe atrophy in early-stage Alzheimer’s disease identified using an improved detection methodology. Psychiatry Res. 155, 11–19. 10.1016/j.pscychresns.2006.12.013

Fernández-Cabello, S., Kronbichler, M., Van Dijk, K.R.A., Goodman, J.A., Spreng, R.N., Schmitz, T.W., on behalf of the Alzheimer’s Disease Neuroimaging Initiative, 2020. Basal forebrain volume reliably predicts the cortical spread of Alzheimer’s degeneration. Brain 143, 993–1009. 10.1093/brain/awaa012

Fischer, W., Gage, F.H., Björklund, A., 1989. Degenerative Changes in Forebrain Cholinergic Nuclei Correlate with Cognitive Impairments in Aged Rats. Eur. J. Neurosci. 1, 34–45. 10.1111/j.1460-9568.1989.tb00772.x

Fox, P.T., Laird, A.R., Fox, S.P., Fox, P.M., Uecker, A.M., Crank, M., Koenig, S.F., Lancaster, J.L., 2005. Brainmap taxonomy of experimental design: Description and evaluation. Hum. Brain Mapp. 25, 185–198. 10.1002/hbm.20141

Fox, P.T., Lancaster, J.L., 2002. Mapping context and content: the BrainMap model. Nat. Rev. Neurosci. 3, 319–321. 10.1038/nrn789

Frisoni, G.B., 2002. Detection of grey matter loss in mild Alzheimer’s disease with voxel based morphometry. J. Neurol. Neurosurg. Psychiatry 73, 657–664. 10.1136/jnnp.73.6.657

Frisoni, G.B., Fox, N.C., Jack, C.R., Scheltens, P., Thompson, P.M., 2010. The clinical use of structural MRI in Alzheimer disease. Nat. Rev. Neurol. 6, 67–77. 10.1038/nrneurol.2009.215

Futami, T., Takakusaki, K., Kitai, S.T., 1995. Glutamatergic and cholinergic inputs from the pedunculopontine tegmental nucleus to dopamine neurons in the substantia nigra pars compacta. Neurosci Res 21, 331–42.

Gais, S., Born, J., 2004. Low acetylcholine during slow-wave sleep is critical for declarative memory consolidation. Proc Natl Acad Sci U A 101, 2140–4.

Gerardin, E., Chételat, G., Chupin, M., Cuingnet, R., Desgranges, B., Kim, H.-S., Niethammer, M., Dubois, B., Lehéricy, S., Garnero, L., Eustache, F., Colliot, O., Alzheimer’s Disease Neuroimaging Initiative, 2009. Multidimensional classification of hippocampal shape features discriminates Alzheimer’s disease and mild cognitive impairment from normal aging. NeuroImage 47, 1476–1486. 10.1016/j.neuroimage.2009.05.036

Geula, C., Dunlop, S.R., Ayala, I., Kawles, A.S., Flanagan, M.E., Gefen, T., Mesulam, M.-M., 2021. Basal forebrain cholinergic system in the dementias: Vulnerability, resilience, and resistance. J. Neurochem. 158, 1394–1411. 10.1111/jnc.15471

Geula, C., Mesulam, M.-M., 1996. Systematic Regional Variations in the Loss of Cortical Cholinergic Fibers in Alzheimer’s Disease. Cereb. Cortex 6, 165–177. 10.1093/cercor/6.2.165

Gili, T., Cercignani, M., Serra, L., Perri, R., Giove, F., Maraviglia, B., Caltagirone, C., Bozzali, M., 2011. Regional brain atrophy and functional disconnection across Alzheimer’s disease evolution. J. Neurol. Neurosurg. Psychiatry 82, 58–66. 10.1136/jnnp.2009.199935

Gold, B.T., Jiang, Y., Jicha, G.A., Smith, C.D., 2010. Functional response in ventral temporal cortex differentiates mild cognitive impairment from normal aging. Hum. Brain Mapp. 31, 1249–1259. 10.1002/hbm.20932

Good, C.D., Scahill, R.I., Fox, N.C., Ashburner, J., Friston, K.J., Chan, D., Crum, W.R., Rossor, M.N., Frackowiak, R.S.J., 2002. Automatic Differentiation of Anatomical Patterns in the Human Brain: Validation with Studies of Degenerative Dementias. NeuroImage 17, 29–46. 10.1006/nimg.2002.1202

Grothe, M., Heinsen, H., Teipel, S., 2013. Longitudinal measures of cholinergic forebrain atrophy in the transition from healthy aging to Alzheimer’s disease. Neurobiol. Aging 34, 1210–1220. 10.1016/j.neurobiolaging.2012.10.018

Grothe, M., Heinsen, H., Teipel, S.J., 2012. Atrophy of the Cholinergic Basal Forebrain Over the Adult Age Range and in Early Stages of Alzheimer’s Disease. Biol. Psychiatry 71, 805–813. 10.1016/j.biopsych.2011.06.019

Guan, X., Guo, T., Zhou, C., Wu, J., Zeng, Q., Li, K., Luo, X., Bai, X., Wu, H., Gao, T., Gu, L., Liu, X., Cao, Z., Wen, J., Chen, J., Wei, H., Zhang, Y., Liu, C., Song, Z., Yan, Y., Pu, J., Zhang, B., Xu, X., Zhang, M., 2022. Altered brain iron depositions from aging to Parkinson’s disease and Alzheimer’s disease: A quantitative susceptibility mapping study. NeuroImage 264, 119683. 10.1016/j.neuroimage.2022.119683

Guitart-Masip, M., Salami, A., Garrett, D., Rieckmann, A., Lindenberger, U., Bäckman, L., 2016. BOLD Variability is Related to Dopaminergic Neurotransmission and Cognitive Aging. Cereb. Cortex 26, 2074–2083. 10.1093/cercor/bhv029

Guo, X., Wang, Z., Li, K., Li, Z., Qi, Z., Jin, Z., Yao, L., Chen, K., 2010. Voxel-based assessment of gray and white matter volumes in Alzheimer’s disease. Neurosci. Lett. 468, 146–150. 10.1016/j.neulet.2009.10.086

Hall, A.M., Moore, R.Y., Lopez, O.L., Kuller, L., Becker, J.T., 2008. Basal forebrain atrophy is a presymptomatic marker for Alzheimer’s disease. Alzheimers Dement. J. Alzheimers Assoc. 4, 271–279. 10.1016/j.jalz.2008.04.005

Hämäläinen, A., Pihlajamäki, M., Tanila, H., Hänninen, T., Niskanen, E., Tervo, S., Karjalainen, P.A., Vanninen, R.L., Soininen, H., 2007. Increased fMRI responses during encoding in mild cognitive impairment. Neurobiol. Aging 28, 1889–1903. 10.1016/j.neurobiolaging.2006.08.008

Hasselmo, M.E., 2006. The role of acetylcholine in learning and memory. Curr. Opin. Neurobiol. 16, 710–715. 10.1016/j.conb.2006.09.002

Hasselmo, M.E., Sarter, M., 2011. Modes and models of forebrain cholinergic neuromodulation of cognition. Neuropsychopharmacology 36, 52–73.

Heckner, M.K., Cieslik, E.C., Küppers, V., Fox, P.T., Eickhoff, S.B., Langner, R., 2021. Delineating visual, auditory and motor regions in the human brain with functional neuroimaging: a BrainMap-based meta-analytic synthesis. Sci. Rep. 11, 9942. 10.1038/s41598-021-88773-9

Hett, K., Ta, V.-T., Catheline, G., Tourdias, T., Manjón, J.V., Coupé, P., Alzheimer’s Disease Neuroimaging Initiative, Weiner, M.W., Aisen, P., Petersen, R., Jack, C.R., Jagust, W., Trojanowki, J.Q., Toga, A.W., Beckett, L., Green, R.C., Saykin, A.J., Morris, J., Shaw, L.M., Khachaturian, Z., Sorensen, G., Carrillo, M., Kuller, L., Raichle, M., Paul, S., Davies, P., Fillit, H., Hefti, F., Holtzman, D., Mesulam, M.M., Potter, W., Snyder, P., Montine, T., Thomas, R.G., Donohue, M., Walter, S., Sather, T., Jiminez, G., Balasubramanian, A.B., Mason, J., Sim, I., Harvey, D., Bernstein, M., Fox, N., Thompson, P., Schuff, N., DeCArli, C., Borowski, B., Gunter, J., Senjem, M., Vemuri, P., Jones, D., Kantarci, K., Ward, C., Koeppe, R.A., Foster, N., Reiman, E.M., Chen, K., Mathis, C., Landau, S., Cairns, N.J., Householder, E., Taylor-Reinwald, L., Lee, V., Korecka, M., Figurski, M., Crawford, K., Neu, S., Foroud, T.M., Potkin, S., Shen, L., Faber, K., Kim, S., Nho, K., Thal, L., Frank, R., Hsiao, J., Kaye, J., Quinn, J., Silbert, L., Lind, B., Carter, R., Dolen, S., Ances, B., Carroll, M., Creech, M.L., Franklin, E., Mintun, M.A., Schneider, S., Oliver, A., Schneider, L.S., Pawluczyk, S., Beccera, M., Teodoro, L., Spann, B.M., Brewer, J., Vanderswag, H., Fleisher, A., Marson, D., Griffith, R., Clark, D., Geldmacher, D., Brockington, J., Roberson, E., Love, M.N., Heidebrink, J.L., Lord, J.L., Mason, S.S., Albers, C.S., Knopman, D., Johnson, Kris, Grossman, H., Mitsis, E., Shah, R.C., deToledo-Morrell, L., Doody, R.S., Villanueva-Meyer, J., Chowdhury, M., Rountree, S., Dang, M., Duara, R., Varon, D., Greig, M.T., Roberts, P., Stern, Y., Honig, L.S., Bell, K.L., Albert, M., Onyike, C., D’Agostino, D., Kielb, S., Galvin, J.E., Cerbone, B., Michel, C.A., Pogorelec, D.M., Rusinek, H., de Leon, M.J., Glodzik, L., De Santi, S., Womack, K., Mathews, D., Quiceno, M., Doraiswamy, P.M., Petrella, J.R., Borges-Neto, S., Wong, T.Z., Coleman, E., Levey, A.I., Lah, J.J., Cella, J.S., Burns, J.M., Swerdlow, R.H., Brooks, W.M., Arnold, S.E., Karlawish, J.H., Wolk, D., Clark, C.M., Apostolova, L., Tingus, K., Woo, E., Silverman, D.H.S., Lu, P.H., Bartzokis, G., Smith, C.D., Jicha, G., Hardy, P., Sinha, P., Oates, E., Conrad, G., Graff-Radford, N.R., Parfitt, F., Kendall, T., Johnson, H., Lopez, O.L., Oakley, M., Simpson, D.M., Farlow, M.R., Hake, A.M., Matthews, B.R., Brosch, J.R., Herring, S., Hunt, C., Porsteinsson, A.P., Goldstein, B.S., Martin, K., Makino, K.M., Ismail, M.S., Brand, C., Mulnard, R.A., Thai, G., Mc-Adams-Ortiz, C., van Dyck, C.H., Carson, R.E., MacAvoy, M.G., Varma, P., Chertkow, H., Bergman, H., Hosein, C., Black, S., Stefanovic, B., Caldwell, C., Hsiung, G.-Y.R., Feldman, H., Mudge, B., Assaly, M., Finger, Elizabeth, Pasternack, S., Rachisky, I., Trost, D., Kertesz, A., Bernick, C., Munic, D., Lipowski, K., Weintraub, M.A.S., Bonakdarpour, B., Kerwin, D., Wu, C.-K., Johnson, N., Sadowsky, C., Villena, T., Turner, R.S., Johnson, Kathleen, Reynolds, B., Sperling, R.A., Johnson, K.A., Marshall, G., Yesavage, J., Taylor, J.L., Lane, B., Rosen, A., Tinklenberg, J., Sabbagh, M.N., Belden, C.M., Jacobson, S.A., Sirrel, S.A., Kowall, N., Killiany, R., Budson, A.E., Norbash, A., Johnson, P.L., Obisesan, T.O., Wolday, S., Allard, J., Lerner, A., Ogrocki, P., Tatsuoka, C., Fatica, P., Fletcher, E., Maillard, P., Olichney, J., Carmichael, O., Kittur, S., Borrie, M., Lee, T.-Y., Bartha, R., Johnson, S., Asthana, S., Carlsson, C.M., Preda, A., Nguyen, D., Tariot, P., Burke, A., Trncic, N., Fleisher, A., Reeder, S., Bates, V., Capote, H., Rainka, M., Scharre, D.W., Kataki, M., Adeli, A., Zimmerman, E.A., Celmins, D., Brown, A.D., Pearlson, G.D., Blank, K., Anderson, K., Flashman, L.A., Seltzer, M., Hynes, M.L., Santulli, R.B., Sink, K.M., Gordineer, L., Williamson, J.D., Garg, P., Watkins, F., Ott, B.R., Querfurth, H., Tremont, G., Salloway, S., Malloy, P., Correia, S., Rosen, H.J., Miller, B.L., Perry, D., Mintzer, J., Spicer, K., Bachman, D., Finger, Elizabether, Pasternak, S., Rachinsky, I., Rogers, J., Drost, D., Pomara, N., Hernando, R., Sarrael, A., Schultz, S.K., Boles Ponto, L.L., Shim, H., Smith, K.E., Relkin, N., Chaing, G., Lin, M., Ravdin, L., Smith, A., Raj, B.A., Fargher, K., 2019. Multimodal Hippocampal Subfield Grading For Alzheimer’s Disease Classification. Sci. Rep. 9, 13845. 10.1038/s41598-019-49970-9

Heys, J.G., Giocomo, L.M., Hasselmo, M.E., 2010. Cholinergic Modulation of the Resonance Properties of Stellate Cells in Layer II of Medial Entorhinal Cortex. J. Neurophysiol. 104, 258–270. 10.1152/jn.00492.2009

Insel, P.S., Mormino, E.C., Aisen, P.S., Thompson, W.K., Donohue, M.C., 2020. Neuroanatomical spread of amyloid β and tau in Alzheimer’s disease: implications for primary prevention. Brain Commun. 2, fcaa007. 10.1093/braincomms/fcaa007

Irish, M., Eyre, N., Dermody, N., O’Callaghan, C., Hodges, J.R., Hornberger, M., Piguet, O., 2016. Neural Substrates of Semantic Prospection - Evidence from the Dementias. Front. Behav. Neurosci. 10, 96. 10.3389/fnbeh.2016.00096

Irish, M., Hodges, J.R., Piguet, O., 2013. Episodic future thinking is impaired in the behavioural variant of frontotemporal dementia. Cortex J. Devoted Study Nerv. Syst. Behav. 49, 2377–2388. 10.1016/j.cortex.2013.03.002

Ishii, K., Sasaki, H., Kono, A.K., Miyamoto, N., Fukuda, T., Mori, E., 2005. Comparison of gray matter and metabolic reduction in mild Alzheimer’s disease using FDG-PET and voxel-based morphometric MR studies. Eur. J. Nucl. Med. Mol. Imaging 32, 959–963. 10.1007/s00259-004-1740-5

Jack, C.R., Bennett, D.A., Blennow, K., Carrillo, M.C., Dunn, B., Haeberlein, S.B., Holtzman, D.M., Jagust, W., Jessen, F., Karlawish, J., Liu, E., Molinuevo, J.L., Montine, T., Phelps, C., Rankin, K.P., Rowe, C.C., Scheltens, P., Siemers, E., Snyder, H.M., Sperling, R., Contributors, Elliott, C., Masliah, E., Ryan, L., Silverberg, N., 2018. NIA-AA Research Framework: Toward a biological definition of Alzheimer’s disease. Alzheimers Dement. 14, 535–562. 10.1016/j.jalz.2018.02.018

Jagust, W., 2018. Imaging the evolution and pathophysiology of Alzheimer disease. Nat. Rev. Neurosci. 19, 687–700. 10.1038/s41583-018-0067-3

Jessen, F., Amariglio, R.E., van Boxtel, M., Breteler, M., Ceccaldi, M., Chételat, G., Dubois, B., Dufouil, C., Ellis, K.A., van der Flier, W.M., Glodzik, L., van Harten, A.C., de Leon, M.J., McHugh, P., Mielke, M.M., Molinuevo, J.L., Mosconi, L., Osorio, R.S., Perrotin, A., Petersen, R.C., Rabin, L.A., Rami, L., Reisberg, B., Rentz, D.M., Sachdev, P.S., de la Sayette, V., Saykin, A.J., Scheltens, P., Shulman, M.B., Slavin, M.J., Sperling, R.A., Stewart, R., Uspenskaya, O., Vellas, B., Visser, P.J., Wagner, M., Group, S.C.D.I. (SCD-I.W., 2014. A conceptual framework for research on subjective cognitive decline in preclinical Alzheimer’s disease. Alzheimers Dement. 10, 844–852. 10.1016/j.jalz.2014.01.001

Jessen, F., Feyen, L., Freymann, K., Tepest, R., Maier, W., Heun, R., Schild, H.-H., Scheef, L., 2006. Volume reduction of the entorhinal cortex in subjective memory impairment. Neurobiol. Aging 27, 1751–1756. 10.1016/j.neurobiolaging.2005.10.010

Johnson, K.A., Fox, N.C., Sperling, R.A., Klunk, W.E., 2012. Brain Imaging in Alzheimer Disease. Cold Spring Harb. Perspect. Med. 2, a006213–a006213. 10.1101/cshperspect.a006213

Kanda, T., Ishii, K., Uemura, T., Miyamoto, N., Yoshikawa, T., Kono, A.K., Mori, E., 2008. Comparison of grey matter and metabolic reductions in frontotemporal dementia using FDG-PET and voxel-based morphometric MR studies. Eur. J. Nucl. Med. Mol. Imaging 35, 2227–2234. 10.1007/s00259-008-0871-5

Karas, G.B., Scheltens, P., Rombouts, S. a. R.B., Visser, P.J., van Schijndel, R.A., Fox, N.C., Barkhof, F., 2004. Global and local gray matter loss in mild cognitive impairment and Alzheimer’s disease. NeuroImage 23, 708–716. 10.1016/j.neuroimage.2004.07.006

Kerbler, G.M., Fripp, J., Rowe, C.C., Villemagne, V.L., Salvado, O., Rose, S., Coulson, E.J., 2015. Basal forebrain atrophy correlates with amyloid β burden in Alzheimer’s disease. NeuroImage Clin. 7, 105–113. 10.1016/j.nicl.2014.11.015

Khan, U.A., Liu, L., Provenzano, F.A., Berman, D.E., Profaci, C.P., Sloan, R., Mayeux, R., Duff, K.E., Small, S.A., 2014. Molecular drivers and cortical spread of lateral entorhinal cortex dysfunction in preclinical Alzheimer’s disease. Nat. Neurosci. 17, 304–311. 10.1038/nn.3606

Kilimann, I., Grothe, M., Heinsen, H., Alho, E.J.L., Grinberg, L., Amaro Jr., E., dos Santos, G.A.B., da Silva, R.E., Mitchell, A.J., Frisoni, G.B., Bokde, A.L.W., Fellgiebel, A., Filippi, M., Hampel, H., Klöppel, S., Teipel, S.J., 2014. Subregional Basal Forebrain Atrophy in Alzheimer’s Disease: A Multicenter Study. J. Alzheimers Dis. 40, 687–700. 10.3233/JAD-132345

Kim, S., Youn, Y.C., Hsiung, G.-Y.R., Ha, S.-Y., Park, K.-Y., Shin, H.-W., Kim, D.-K., Kim, S.-S., Kee, B.S., 2011. Voxel-based morphometric study of brain volume changes in patients with Alzheimer’s disease assessed according to the Clinical Dementia Rating score. J. Clin. Neurosci. Off. J. Neurosurg. Soc. Australas. 18, 916–921. 10.1016/j.jocn.2010.12.019

Kukolja, J., Thiel, C.M., Fink, G.R., 2009. Cholinergic Stimulation Enhances Neural Activity Associated with Encoding but Reduces Neural Activity Associated with Retrieval in Humans. J. Neurosci. 29, 8119–8128. 10.1523/JNEUROSCI.0203-09.2009

Laird, A.R., Eickhoff, S.B., Kurth, F., Fox, P.M., Uecker, A.M., Turner, J.A., Robinson, J.L., Lancaster, J.L., Fox, P.T., 2009. ALE Meta-Analysis Workflows Via the Brainmap Database: Progress Towards A Probabilistic Functional Brain Atlas. Front. Neuroinformatics 3, 23. 10.3389/neuro.11.023.2009

Laird, A.R., Fox, P.M., Price, C.J., Glahn, D.C., Uecker, A.M., Lancaster, J.L., Turkeltaub, P.E., Kochunov, P., Fox, P.T., 2005. ALE meta-analysis: controlling the false discovery rate and performing statistical contrasts. Hum. Brain Mapp. 25, 155–164. 10.1002/hbm.20136

Laird, A.R., Robinson, J.L., McMillan, K.M., Tordesillas-Gutiérrez, D., Moran, S.T., Gonzales, S.M., Ray, K.L., Franklin, C., Glahn, D.C., Fox, P.T., Lancaster, J.L., 2010. Comparison of the disparity between Talairach and MNI coordinates in functional neuroimaging data: Validation of the Lancaster transform. NeuroImage 51, 677–683. 10.1016/j.neuroimage.2010.02.048

Lancaster, J.L., Tordesillas-Gutiérrez, D., Martinez, M., Salinas, F., Evans, A., Zilles, K., Mazziotta, J.C., Fox, P.T., 2007. Bias between MNI and Talairach coordinates analyzed using the ICBM-152 brain template. Hum. Brain Mapp. 28, 1194–1205. 10.1002/hbm.20345

LeDoux, J., 2003. The emotional brain, fear, and the amygdala. Cell Mol Neurobiol 23, 727–38.

Leutritz, T., Seif, M., Helms, G., Samson, R.S., Curt, A., Freund, P., Weiskopf, N., 2020. Multiparameter mapping of relaxation (R1, R2*), proton density and magnetization transfer saturation at 3 T: A multicenter dual-vendor reproducibility and repeatability study. Hum. Brain Mapp. 41, 4232–4247. 10.1002/hbm.25122

Li, X., Yu, B., Sun, Q., Zhang, Y., Ren, M., Zhang, X., Li, A., Yuan, J., Madisen, L., Luo, Q., Zeng, H., Gong, H., Qiu, Z., 2018. Generation of a whole-brain atlas for the cholinergic system and mesoscopic projectome analysis of basal forebrain cholinergic neurons. Proc. Natl. Acad. Sci. 115, 415–420. 10.1073/pnas.1703601115

Lisman, J., Grace, A.A., Duzel, E., 2011. A neoHebbian framework for episodic memory; role of dopamine-dependent late LTP. Trends Neurosci 34, 536–47.

Lisman, J.E., Grace, A.A., 2005. The Hippocampal-VTA Loop: Controlling the Entry of Information into Long-Term Memory. Neuron 46, 703–13.

Liu, A.K.L., Chang, R.C.-C., Pearce, R.K.B., Gentleman, S.M., 2015. Nucleus basalis of Meynert revisited: anatomy, history and differential involvement in Alzheimer’s and Parkinson’s disease. Acta Neuropathol. (Berl.) 129, 527–540. 10.1007/s00401-015-1392-5

Liu, L., Drouet, V., Wu, J.W., Witter, M.P., Small, S.A., Clelland, C., Duff, K., 2012. Trans-Synaptic Spread of Tau Pathology In Vivo. PLoS ONE 7, e31302. 10.1371/journal.pone.0031302

Loeffler, D.A., Connor, J.R., Juneau, P.L., Snyder, B.S., Kanaley, L., DeMaggio, A.J., Nguyen, H., Brickman, C.M., LeWitt, P.A., 1995. Transferrin and Iron in Normal, Alzheimer’s Disease, and Parkinson’s Disease Brain Regions. J. Neurochem. 65, 710–716. 10.1046/j.1471-4159.1995.65020710.x

Long, Z., Jing, B., Yan, H., Dong, J., Liu, H., Mo, X., Han, Y., Li, H., 2016. A support vector machine-based method to identify mild cognitive impairment with multi-level characteristics of magnetic resonance imaging. Neuroscience 331, 169–176. 10.1016/j.neuroscience.2016.06.025

Mark, G.P., Shabani, S., Dobbs, L.K., Hansen, S.T., 2011. Cholinergic modulation of mesolimbic dopamine function and reward. Physiol. Behav. 104, 76–81. 10.1016/j.physbeh.2011.04.052

Matsuda, H., Kitayama, N., Ohnishi, T., Asada, T., Nakano, S., Sakamoto, S., Imabayashi, E., Katoh, A., 2002. Longitudinal evaluation of both morphologic and functional changes in the same individuals with Alzheimer’s disease. J. Nucl. Med. Off. Publ. Soc. Nucl. Med. 43, 304–311.

Mechelli, A., Price, C., Friston, K., Ashburner, J., 2005. Voxel-Based Morphometry of the Human Brain: Methods and Applications. Curr. Med. Imaging Rev. 1, 105–113. 10.2174/1573405054038726

Mesulam, 2004. The cholinergic innervation of the human cerebral cortex. Prog. Brain Res. 145, 67–78. 10.1016/S0079-6123(03)45004-8

Mesulam, Geula, 1988. Nucleus basalis (Ch4) and cortical cholinergic innervation in the human brain: Observations based on the distribution of acetylcholinesterase and choline acetyltransferase. J. Comp. Neurol. 275, 216–240. 10.1002/cne.902750205

Mesulam, M., 2004. The Cholinergic Lesion of Alzheimer’s Disease: Pivotal Factor or Side Show? Learn. Mem. 11, 43–49. 10.1101/lm.69204

Mesulam, M., Shaw, P., Mash, D., Weintraub, S., 2004. Cholinergic nucleus basalis tauopathy emerges early in the aging-MCI-AD continuum. Ann. Neurol. 55, 815–828. 10.1002/ana.20100

Mesulam, M.-M., Hersh, L.B., Mash, D.C., Geula, C., 1992. Differential cholinergic innervation within functional subdivisions of the human cerebral cortex: A choline acetyltransferase study. J. Comp. Neurol. 318, 316–328. 10.1002/cne.903180308

Mesulam, M.-M., Mufson, E.J., Levey, A.I., Wainer, B.H., 1983. Cholinergic innervation of cortex by the basal forebrain: Cytochemistry and cortical connections of the septal area, diagonal band nuclei, nucleus basalis (Substantia innominata), and hypothalamus in the rhesus monkey. J. Comp. Neurol. 214, 170–197. 10.1002/cne.902140206

Miller, A.D., Blaha, C.D., 2004. Nigrostriatal dopamine release modulated by mesopontine muscarinic receptors. Neuroreport 15, 1805–8.

Mitsushima, D., Sano, A., Takahashi, T., 2013. A cholinergic trigger drives learning-induced plasticity at hippocampal synapses. Nat. Commun. 4. 10.1038/ncomms3760

Müller, Cieslik, E.C., Laird, A.R., Fox, P.T., Radua, J., Mataix-Cols, D., Tench, C.R., Yarkoni, T., Nichols, T.E., Turkeltaub, P.E., Wager, T.D., Eickhoff, S.B., 2018. Ten simple rules for neuroimaging meta-analysis. Neurosci. Biobehav. Rev. 84, 151–161. 10.1016/j.neubiorev.2017.11.012

Muth, K., Schönmeyer, R., Matura, S., Haenschel, C., Schröder, J., Pantel, J., 2010. Mild Cognitive Impairment in the Elderly is Associated with Volume Loss of the Cholinergic Basal Forebrain Region. Biol. Psychiatry 67, 588–591. 10.1016/j.biopsych.2009.02.026

Oakman, S.A., Faris, P.L., Kerr, P.E., Cozzari, C., Hartman, B.K., 1995. Distribution of pontomesencephalic cholinergic neurons projecting to substantia nigra differs significantly from those projecting to ventral tegmental area. J Neurosci 15, 5859–69.

Ohnishi, T., Matsuda, H., Tabira, T., Asada, T., Uno, M., 2001. Changes in brain morphology in Alzheimer disease and normal aging: is Alzheimer disease an exaggerated aging process? AJNR Am. J. Neuroradiol. 22, 1680–1685.

Ostrerova-Golts, N., Petrucelli, L., Hardy, J., Lee, J.M., Farer, M., Wolozin, B., 2000. The A53T α-Synuclein Mutation Increases Iron-Dependent Aggregation and Toxicity. J. Neurosci. 20, 6048–6054. 10.1523/JNEUROSCI.20-16-06048.2000

Page, M.J., McKenzie, J.E., Bossuyt, P.M., Boutron, I., Hoffmann, T.C., Mulrow, C.D., Shamseer, L., Tetzlaff, J.M., Akl, E.A., Brennan, S.E., Chou, R., Glanville, J., Grimshaw, J.M., Hróbjartsson, A., Lalu, M.M., Li, T., Loder, E.W., Mayo-Wilson, E., McDonald, S., McGuinness, L.A., Stewart, L.A., Thomas, J., Tricco, A.C., Welch, V.A., Whiting, P., Moher, D., 2021. The PRISMA 2020 statement: an updated guideline for reporting systematic reviews. BMJ n71. 10.1136/bmj.n71

Pennanen, C., Testa, C., Laakso, M.P., Hallikainen, M., Helkala, E.-L., Hänninen, T., 2005. A voxel based morphometry study on mild cognitive impairment. J. Neurol. Neurosurg. Psychiatry 76, 11–14. 10.1136/jnnp.2004.035600

Petersen, R.C., Lopez, O., Armstrong, M.J., Getchius, T.S.D., Ganguli, M., Gloss, D., Gronseth, G.S., Marson, D., Pringsheim, T., Day, G.S., Sager, M., Stevens, J., Rae-Grant, A., 2018. Practice guideline update summary: Mild cognitive impairment: Report of the Guideline Development, Dissemination, and Implementation Subcommittee of the American Academy of Neurology. Neurology 90, 126–135. 10.1212/WNL.0000000000004826

Picciotto, M.R., Higley, M.J., Mineur, Y.S., 2012. Acetylcholine as a Neuromodulator: Cholinergic Signaling Shapes Nervous System Function and Behavior. Neuron 76, 116–129. 10.1016/j.neuron.2012.08.036

Planche, V., Manjon, J.V., Mansencal, B., Lanuza, E., Tourdias, T., Catheline, G., Coupé, P., 2022. Structural progression of Alzheimer’s disease over decades: the MRI staging scheme. Brain Commun. 4, fcac109. 10.1093/braincomms/fcac109

Poulin, S.P., Dautoff, R., Morris, J.C., Barrett, L.F., Dickerson, B.C., 2011. Amygdala atrophy is prominent in early Alzheimer’s disease and relates to symptom severity. Psychiatry Res. 194, 7–13. 10.1016/j.pscychresns.2011.06.014

Prieto del Val, L., Cantero, J.L., Atienza, M., 2016. Atrophy of amygdala and abnormal memory-related alpha oscillations over posterior cingulate predict conversion to Alzheimer’s disease. Sci. Rep. 6, 31859. 10.1038/srep31859

Rajmohan, R., Reddy, P.H., 2017. Amyloid Beta and Phosphorylated Tau Accumulations Cause Abnormalities at Synapses of Alzheimer’s disease Neurons. J. Alzheimers Dis. JAD 57, 975–999. 10.3233/JAD-160612

Rami, L., Gómez-Anson, B., Monte, G.C., Bosch, B., Sánchez-Valle, R., Molinuevo, J.L., 2009. Voxel based morphometry features and follow-up of amnestic patients at high risk for Alzheimer’s disease conversion. Int. J. Geriatr. Psychiatry 24, 875–884. 10.1002/gps.2216

Ramos Bernardes da Silva Filho, S., Oliveira Barbosa, J.H., Rondinoni, C., dos Santos, A.C., Garrido Salmon, C.E., da Costa Lima, N.K., Ferriolli, E., Moriguti, J.C., 2017. Neurodegeneration profile of Alzheimer’s patients: A brain morphometry study. NeuroImage Clin. 15, 15–24. 10.1016/j.nicl.2017.04.001

Rémy, F., Mirrashed, F., Campbell, B., Richter, W., 2005. Verbal episodic memory impairment in Alzheimer’s disease: a combined structural and functional MRI study. NeuroImage 25, 253–266. 10.1016/j.neuroimage.2004.10.045

Rogers, J.T., Bush, A.I., Cho, H.-H., Smith, D.H., Thomson, A.M., Friedrlich, A.L., Lahiri, D.K., Leedman, P.J., Huang, X., Cahill, C.M., 2008. Iron and the translation of the amyloid precursor protein (APP) and ferritin mRNAs: Riboregulation against neural oxidative damage in Alzheimer’s disease. Biochem. Soc. Trans. 36, 1282–1287. 10.1042/BST0361282

Rutishauser, U., Ross, I.B., Mamelak, A.N., Schuman, E.M., 2010. Human memory strength is predicted by theta-frequency phase-locking of single neurons. Nature 464, 903–907. 10.1038/nature08860

Schliebs, R., Arendt, T., 2011. The cholinergic system in aging and neuronal degeneration. Behav. Brain Res. 221, 555–563. 10.1016/j.bbr.2010.11.058

Schmitz, T.W., Duncan, J., 2018. Normalization and the Cholinergic Microcircuit: A Unified Basis for Attention. Trends Cogn. Sci. 22, 422–437. 10.1016/j.tics.2018.02.011

Schmitz, T.W., Mur, M., Aghourian, M., Bedard, M.-A., Spreng, R.N., 2018. Longitudinal Alzheimer’s Degeneration Reflects the Spatial Topography of Cholinergic Basal Forebrain Projections. Cell Rep. 24, 38–46. 10.1016/j.celrep.2018.06.001

Schmitz, T.W., Soreq, H., Poirier, J., Spreng, R.N., 2020. Longitudinal Basal Forebrain Degeneration Interacts with TREM2/C3 Biomarkers of Inflammation in Presymptomatic Alzheimer’s Disease. J. Neurosci. 40, 1931–1942. 10.1523/JNEUROSCI.1184-19.2019

Schmitz, T.W., Spreng, R.N., 2016. Basal forebrain degeneration precedes and predicts the cortical spread of Alzheimer’s pathology. Nat. Commun. 7, 13249. 10.1038/ncomms13249

Sherman, S.J., Atri, A., Hasselmo, M.E., Stern, C.E., Howard, M.W., 2003. Scopolamine impairs human recognition memory: data and modeling. Behav Neurosci 117, 526–39.

Shiino, A., Watanabe, T., Maeda, K., Kotani, E., Akiguchi, I., Matsuda, M., 2006. Four subgroups of Alzheimer’s disease based on patterns of atrophy using VBM and a unique pattern for early onset disease. NeuroImage 33, 17–26. 10.1016/j.neuroimage.2006.06.010

Sian-Hülsmann, J., Mandel, S., Youdim, M.B.H., Riederer, P., 2011. The relevance of iron in the pathogenesis of Parkinson’s disease. J. Neurochem. 118, 939–957. 10.1111/j.1471-4159.2010.07132.x

Sperling, R., 2011. The potential of functional MRI as a biomarker in early Alzheimer’s disease. Neurobiol. Aging 32, S37–S43. 10.1016/j.neurobiolaging.2011.09.009

Steiger, T.K., Bunzeck, N., 2017. Reward Dependent Invigoration Relates to Theta Oscillations and Is Predicted by Dopaminergic Midbrain Integrity in Healthy Elderly. Front. Aging Neurosci. 9. 10.3389/fnagi.2017.00001

Tardif, C.L., Collins, D.L., Pike, G.B., 2009. Regional impact of field strength on voxel-based morphometry results. Hum. Brain Mapp. 31, 943–957. 10.1002/hbm.20908

Threlfell, S., Lalic, T., Platt, N.J., Jennings, K.A., Deisseroth, K., Cragg, S.J., 2012. Striatal Dopamine Release Is Triggered by Synchronized Activity in Cholinergic Interneurons. Neuron 75, 58–64. 10.1016/j.neuron.2012.04.038

Turkeltaub, P.E., Eickhoff, S.B., Laird, A.R., Fox, M., Wiener, M., Fox, P., 2012. Minimizing within-experiment and within-group effects in activation likelihood estimation meta-analyses. Hum. Brain Mapp. 33, 1–13. 10.1002/hbm.21186

Vanasse, T.J., Fox, P.M., Barron, D.S., Robertson, M., Eickhoff, S.B., Lancaster, J.L., Fox, P.T., 2018. BrainMap VBM: An environment for structural meta-analysis. Hum. Brain Mapp. 39, 3308–3325. 10.1002/hbm.24078

Vogels, O.J.M., Broere, C.A.J., Ter Laak, H.J., Ten Donkelaar, H.J., Nieuwenhuys, R., Schulte, B.P.M., 1990. Cell loss and shrinkage in the nucleus basalis Meynert complex in Alzheimer’s disease. Neurobiol. Aging 11, 3–13. 10.1016/0197-4580(90)90056-6

Ward, R.J., Zucca, F.A., Duyn, J.H., Crichton, R.R., Zecca, L., 2014. The role of iron in brain ageing and neurodegenerative disorders. Lancet Neurol. 13, 1045–1060. 10.1016/S1474-4422(14)70117-6

Warren, S.L., Moustafa, A.A., 2023. Functional magnetic resonance imaging, deep learning, and Alzheimer’s disease: A systematic review. J. Neuroimaging 33, 5–18. 10.1111/jon.13063

Wei, G., Irish, M., Hodges, J.R., Piguet, O., Kumfor, F., 2020. Disease-specific profiles of apathy in Alzheimer’s disease and behavioural-variant frontotemporal dementia differ across the disease course. J. Neurol. 267, 1086–1096. 10.1007/s00415-019-09679-1

Whitehouse, P.J., Price, D.L., Clark, A.W., Coyle, J.T., DeLong, M.R., 1981. Alzheimer disease: Evidence for selective loss of cholinergic neurons in the nucleus basalis. Ann. Neurol. 10, 122–126. 10.1002/ana.410100203

Whitwell, J.L., 2009. Voxel-Based Morphometry: An Automated Technique for Assessing Structural Changes in the Brain. J. Neurosci. 29, 9661–9664. 10.1523/JNEUROSCI.2160-09.2009

Whitwell, J.L., Jack, C.R., Kantarci, K., Weigand, S.D., Boeve, B.F., Knopman, D.S., Drubach, D.A., Tang-Wai, D.F., Petersen, R.C., Josephs, K.A., 2007. Imaging correlates of posterior cortical atrophy. Neurobiol. Aging 28, 1051–1061. 10.1016/j.neurobiolaging.2006.05.026

Wilson, F.A., Rolls, E.T., 1990. Neuronal responses related to the novelty and familarity of visual stimuli in the substantia innominata, diagonal band of Broca and periventricular region of the primate basal forebrain. Exp Brain Res 80, 104–20.

Winters, B.D., Bussey, T.J., 2005. Removal of cholinergic input to perirhinal cortex disrupts object recognition but not spatial working memory in the rat. Eur J Neurosci 21, 2263– 70.

Winters, B.D., Saksida, L.M., Bussey, T.J., 2006. Paradoxical facilitation of object recognition memory after infusion of scopolamine into perirhinal cortex: implications for cholinergic system function. J Neurosci 26, 9520–9.

Wolz, R., Julkunen, V., Koikkalainen, J., Niskanen, E., Zhang, D.P., Rueckert, D., Soininen, H., Lötjönen, J., the Alzheimer’s Disease Neuroimaging Initiative, 2011. Multi-Method Analysis of MRI Images in Early Diagnostics of Alzheimer’s Disease. PLoS ONE 6, e25446. 10.1371/journal.pone.0025446

Wu, H., Williams, J., Nathans, J., 2014. Complete morphologies of basal forebrain cholinergic neurons in the mouse. eLife 3, e02444. 10.7554/eLife.02444

Wu, J.W., Hussaini, S.A., Bastille, I.M., Rodriguez, G.A., Mrejeru, A., Rilett, K., Sanders, D.W., Cook, C., Fu, H., Boonen, R.A.C.M., Herman, M., Nahmani, E., Emrani, S., Figueroa, Y.H., Diamond, M.I., Clelland, C.L., Wray, S., Duff, K.E., 2016. Neuronal activity enhances tau propagation and tau pathology in vivo. Nat. Neurosci. 19, 1085–1092. 10.1038/nn.4328

Xie, S., Xiao, J.X., Gong, G.L., Zang, Y.F., Wang, Y.H., Wu, H.K., Jiang, X.X., 2006. Voxel-based detection of white matter abnormalities in mild Alzheimer disease. Neurology 66, 1845–1849. 10.1212/01.wnl.0000219625.77625.aa

Yamamoto, A., Shin, R.-W., Hasegawa, K., Naiki, H., Sato, H., Yoshimasu, F., Kitamoto, T., 2002. Iron (III) induces aggregation of hyperphosphorylated τ and its reduction to iron (II) reverses the aggregation: Implications in the formation of neurofibrillary tangles of Alzheimer’s disease. J. Neurochem. 82, 1137–1147. 10.1046/j.1471-4159.2002.01061.x

Yousuf, M., Packard, P.A., Fuentemilla, L., Bunzeck, N., 2021. Functional coupling between CA3 and laterobasal amygdala supports schema dependent memory formation. NeuroImage 244, 118563. 10.1016/j.neuroimage.2021.118563

Záborszky, L., Gombkoto, P., Varsanyi, P., Gielow, M.R., Poe, G., Role, L.W., Ananth, M., Rajebhosale, P., Talmage, D.A., Hasselmo, M.E., Dannenberg, H., Minces, V.H., Chiba, A.A., 2018. Specific Basal Forebrain–Cortical Cholinergic Circuits Coordinate Cognitive Operations. J. Neurosci. 38, 9446–9458. 10.1523/JNEUROSCI.1676-18.2018

Zaborszky, L., Hoemke, L., Mohlberg, H., Schleicher, A., Amunts, K., Zilles, K., 2008. Stereotaxic probabilistic maps of the magnocellular cell groups in human basal forebrain. NeuroImage 42, 1127–1141. 10.1016/j.neuroimage.2008.05.055

Zeng, Q., Qiu, T., Li, K., Luo, X., Wang, S., Xu, X., Liu, X., Hong, L., Li, J., Huang, P., Zhang, M., 2022. Increased functional connectivity between nucleus basalis of Meynert and amygdala in cognitively intact elderly along the Alzheimer’s continuum. NeuroImage Clin. 36, 103256. 10.1016/j.nicl.2022.103256

