## Supplementary Material for "Structural Degeneration of the Nucleus basalis of Meynert in Mild Cognitive Impairment and Alzheimer’s Disease – Evidence from an MRI-based Meta-Analysis"

**Tables S1**: Checklist for Neuroimaging Meta-Analyses after Müller et al. (2018).

| | |
|---|---|
| The research question was specifically defined | YES, and it includes the following contrasts:<br>1) CN>MCI<br>2) CN>AD |
| The literature search was systematic | YES, it included the following search criteria in the BrainMap database:<br><br>CN>MCI ALE meta-analysis<br>1) Experiments contrast *is* gray matter<br>2) Experiments context *is* disease<br>3) Experiments observed changes *is* control>patients<br>4) Subjects diagnosis *is* mild cognitive impairment<br>5) Subjects diagnosis *is* not dementia<br><br>CN>AD ALE meta-analysis<br>1) Experiments contrast *is* gray matter<br>2) Experiments context *is* disease<br>3) Experiments observed changes *is* control>patients<br>4) Subjects diagnosis *is* Alzheimer's disease<br>5) Subjects diagnosis *is* not frontotemporal dementia<br>6) Subjects diagnosis *is not* Lewy body dementia<br>7) Subjects diagnosis *is not* Parkinson's disease<br>8) Subjects diagnosis *is not* behavioral variant frontotemporal dementia<br>9) Subjects diagnosis *is not* non-aphasic frontotemporal dementia |
| Detailed inclusion and exclusion criteria were applied | YES, see Table 1 for more information. |
| Sample overlap was taken into account | YES, experiments were carefully checked regarding the inclusion of possible analysis including subgroups of MCI or AD. |
| All experiments use the same search overage (state how brain coverage is assessed and how small volume corrections and conjunctions are taken into account) | YES, the search coverage was the following:<br>- Only whole-brain coverage<br>- ROI studies were excluded |
| Studies are converted to a common reference space | YES, using the following conversion:<br>- Sleuth's implemented automated tool of using the icbm2tal conversion (converting reported Talairach coordinates to MNI coordinates). |
| The study protocol and all analyses was planned beforehand, including the methods and | YES, |

| | |
|---|---|
| parameters used for inference, correction for multiple testing, etc. | 1) Any non-planned analyses are clearly stated as post-hoc in the paper.<br>2) The meta-analysis used the default methods and parameters of the software. |
| The meta-analysis includes diagnostics | YES, the following:<br>Mild cognitive impairment and Alzheimer's disease |

**Table S2:** Overview of included studies for the contrast HC > MCI. Abbreviations: aMCI: amnestic type mild cognitive impairment; HC: healthy controls.

| Title, authors & date of publication | Groups, mean age, sample size | Included experiment / contrast | Statistical threshold and thresholding method | MRI field strength, reference space, analysis software |
|---|---|---|---|---|
| *White Matter Damage in Alzheimer Disease and Its Relationship to Gray Matter Atrophy* (Agosta et al., 2011) | aMCI, 70 years, n = 15<br><br>HC, 70 years, n = 15 | HC > aMCI | $p < 0.001$<br><br>Cluster-wise corrected for multiple comparisons | 1.5 T<br><br>MNI<br><br>SPM5 |
| *Default-mode network activity distinguishes amnestic type mild cognitive impairment from healthy aging: A combined structural and resting-state functional MRI study* (Bai et al., 2008) | aMCI, 71 years, n = 20<br><br>HC, 69 years, n = 20 | HC > aMCI | $p < 0.05$<br><br>Voxel-wise corrected for multiple comparisons | 1.5 T<br><br>Talairach<br><br>SPM5 |
| *Profile of memory impairment and gray matter loss in amnestic mild cognitive impairment* (Barbeau et al., 2008) | aMCI unimpaired in a visual recognition memory test (DMS48), 72 years, n = 12<br><br>aMCI impaired in a visual recognition memory test (DMS48), 67 years, n = 16<br><br>HC, 63 years, n = 28 | HC > aMCI (unimpaired & impaired performance) | $p < 0.05$<br><br>FDR corrected for multiple comparisons | 1.5 T<br><br>Talairach<br><br>SPM2 |

| | | | | |
|---|---|---|---|---|
| *Differential cortical atrophy in subgroups of mild cognitive impairment* (Bell-McGinty et al., 2005) | aMCI converters to AD during follow-up, 76 years, n = 4<br><br>aMCI non-converters during follow-up, 72 years, n = 5<br><br>MCI-MCD (multiple cognitive domain) converters during follow up, 71 years, n = 10<br><br>MCI-MCD (multiple cognitive domain) non-converters during follow up (ICD: G31.84),72 years, n = 18<br><br>HC, 67 years, n = 47 | HC > MCI (MCI-A converters, MCI-A non-converters, MCI-MCD converters, MCI-MCD non-converters) | $p < 0.001$ | 1.5 T<br><br>MNI<br><br>SPM99 |
| *The contribution of voxel-based morphometry in staging patients with mild cognitive impairment* (Bozzali et al., 2006) | MCI converters to AD, 71 years, n = 14<br><br>HC, 66 years, n = 20 | HC > MCI converters | $p < 0.001$ | 1.5 T<br><br>MNI<br><br>SPM2 |
| *The contribution of voxel-based morphometry in staging patients with mild cognitive impairment* (Bozzali et al., 2006) | MCI non-converters to AD, 71 years, n = 8<br><br>HC, 66 years, n = 20 | HC > MCI non-converters | $p < 0.001$ | 1.5 T<br><br>MNI<br><br>SPM2 |
| *Cerebral perfusion correlates of conversion to Alzheimer's disease in amnestic mild cognitive impairment* | Non-converters aMCI, 71 years, n = 14<br><br>HC, 69 years, n = 17 | HC > Non-converters aMCI to AD | $p < 0.001$ | 1.0 T<br><br>MNI |

| | | | | |
|---|---|---|---|---|
| (Caroli et al., 2007) | | | | SPM2 |
| *Mapping gray matter loss with voxel-based morphometry in mild cognitive impairment* (Chételat et al., 2002) | MCI, 71 years, n = 22<br><br>HC, 67 years, n = 22 | HC > MCI | $p < 0.01$<br><br>Corrected for multiple comparisons | 1.5 T<br><br>Talairach<br><br>SPM99 |
| *Functional response in ventral temporal cortex differentiates mild cognitive impairment from normal aging* (Gold et al., 2010) | MCI, 78 years, n = 12<br><br>HC, 77 years, n = 14 | HC > MCI | $p < 0.001$ | 3.0 T<br><br>MNI<br><br>SPM5 |
| *Increased fMRI responses during encoding in mild cognitive impairment* (Hämäläinen et al., 2007) | MCI, 72 years, n = 14<br><br>HC, 71 years, n = 21 | HC > MCI | $p < 0.05$<br><br>Cluster-wise corrected for multiple comparisons | 1.5 T<br><br>MNI<br><br>SPM2 |

| | | | | |
|---|---|---|---|---|
| *A support vector machine-based method to identify mild cognitive impairment with multi-level characteristics of magnetic resonance imaging* (Long et al., 2016) | MCI, 66 years, n = 29<br><br>HC, 62 years, n = 33 | HC > MCI | $p < 0.01$<br><br>Voxel-wise corrected for multiple comparisons | 3.0 T<br><br>MNI<br><br>SPM8 |
| *A voxel-based morphometry study on mild cognitive impairment* (Pennanen et al., 2005) | MCI, 72 years, n = 51<br><br>HC, 74 years, n = 32 | HC > MCI | $p < 0.001$ | 1.5 T<br><br>MNI<br><br>SPM99 |
| *Voxel based morphometry features and follow-up of amnestic patients at high risk for Alzheimer's disease conversion* (Rami et al., 2009) | aMCI, 73 years, n = 14<br><br>HC, 74 years, n = 27 | HC > aMCI | $p < 0.001$ | 1.5 T<br><br>MNI<br><br>SPM2 |
| *Memory performance correlates with gray matter density in the ento-/perirhinal cortex and posterior hippocampus in patients with mild cognitive impairment and healthy controls — A voxel based morphometry study* (Schmidt-Wilcke et al., 2009) | MCI, 66 years, n = 18<br><br>HC, 63 years, n = 18 | HC > MCI | $p < 0.05$<br><br>FWE corrected for multiple comparisons | 1.5 T<br><br>MNI<br><br>SPM2 |
| *Four subgroups of Alzheimer's disease based on patterns of atrophy using VBM and a* | MCI, 68 years, n = 20 | HC > MCI | $p < 0.05$ | 1.5 T |

| | | | | |
|---|---|---|---|---|
| *unique pattern for early-onset disease* (Shiino et al., 2006) | HC, 69 years, n = 88 | | Voxel-wise corrected for multiple comparisons | Talairach<br><br>MedX |
| *Early morphological brain abnormalities in patients with amnestic mild cognitive impairment* (Yin et al., 2014) | aMCI, 67 years, n = 11<br><br>HC, 62 years, n = 22 | HC > aMCI | $p < 0.05$<br><br>Cluster-wise corrected for multiple comparisons | 3.0 T<br><br>MNI<br><br>SPM5 |
| *Selective Changes of Resting-State Brain Oscillations in aMCI: An fMRI Study Using ALFF* (Zhao et al., 2014) | aMCI, 65 years, n = 20<br><br>HC, 67 years, n = 18 | HC > aMCI | $p < 0.01$<br><br>Voxel-wise corrected for multiple comparisons | 3.0 T<br><br>MNI<br>SPM5 |

**Table S3:** Overview of included studies for the experimental condition HC > AD. Abbreviations: aMCI: amnestic type mild cognitive impairment; HC: healthy controls.

| Title, authors & date of publication | Groups, mean age, sample size | Included experiments | Statistical threshold and thresholding method | MRI field strength, reference space, analysis software |
|---|---|---|---|---|
| *In vivo mapping of gray matter loss with voxel-based morphometry in mild Alzheimer's disease* (Baron et al., 2001) | AD, 73 years, n = 19<br><br>HC, 66 years, n = 16 | HC > AD | p < 0.001 | 1.5 T<br><br>Talairach<br><br>SPM2 |
| *Relationship of cognitive measures and gray and white matter in Alzheimer's disease* (Baxter et al., 2006) | AD, 76 years, n = 15<br><br>HC, 76 years, n = 15 | HC > AD | p < 0.0001 | 1.5 T<br><br>MNI<br><br>SPM2 |
| *Cinguloparietal atrophy distinguishes Alzheimer disease from semantic dementia* (Boxer et al., 2003) | AD, 70 years, n = 11<br><br>HC, 65 years, n = 15 | HC > AD | p < 0.05<br><br>Voxel-wise corrected for multiple comparisons | 1.5 T<br><br>MNI<br><br>SPM99 |
| *The contribution of voxel-based morphometry in staging patients with mild cognitive impairment* (Bozzali et al., 2006) | AD, 68 years, n = 22<br><br>HC, 66 years, n = 20 | HC > AD | p < 0.05<br><br>Voxel-wise corrected for multiple comparisons | 1.5 T<br><br>MNI<br><br>SPM2 |
| *Damage to the cingulum contributes to Alzheimer's disease pathophysiology by* | Probable AD, 73 years, n = 31<br><br>HC, 68 years, n = 14 | HC > AD | p < 0.05 | 3.0 T<br><br>MNI |

| | | | | |
|---|---|---|---|---|
| *deafferentation mechanism* (Bozzali et al., 2012) | | | Voxel-wise corrected for multiple comparisons | SPM8 |
| *Basal forebrain atrophy is a distinctive pattern in dementia with Lewy bodies* (Brenneis et al., 2004) | AD, 73 years, n = 10<br><br>HC, 65 years, n = 10 | HC > AD | $p < 0.05$<br><br>Corrected or multiple comparisons | 1.5 T<br><br>MNI<br><br>SPM99 |
| *Patterns of cerebellar volume loss in dementia with Lewy bodies and Alzheimer's disease: A VBM-DARTEL study* (Colloby et al., 2014) | AD, 79 years, n = 47<br><br>HC, 77 years, n = 39 | HC > AD | $p < 0.05$<br><br>Voxel-wise corrected for multiple comparisons | 3.0 T<br><br>MNI<br><br>SPM8 |
| *Simultaneous arterial spin labeling cerebral blood flow and morphological assessments for detection of Alzheimer's disease* (Dashjamts et al., 2011) | AD, 75 years, n = 23<br><br>HC, 73 years, n = 23 | HC > AD | $p < 0.001$ | 3.0 T<br><br>MNI<br><br>SPM8 |
| *Prospective Memory Impairments in Alzheimer's Disease and Behavioral Variant Frontotemporal Dementia: Clinical and Neural Correlates* (Dermody et al., 2016a) | AD, 63 years, n = 12<br><br>HC, 69 years, n = 12 | HC > AD | $p < 0.05$<br><br>Voxel-wise corrected for multiple comparisons | 3.0 T<br><br>MNI<br><br>FSL |
| *Uncovering the Neural Bases of Cognitive and Affective Empathy Deficits in Alzheimer's Disease and the Behavioral-Variant of Frontotemporal Dementia* (Dermody et al., 2016b) | AD, 66 years, n = 25<br><br>HC, 68 years, n = 22 | HC > AD | $p < 0.05$<br><br>Voxel-wise corrected for | 3.0 T<br><br>MNI<br><br>FSL |

| | | | | |
|---|---|---|---|---|
| | | | multiple comparisons | |
| *Episodic memory impairment in patients with Alzheimer's disease is correlated with entorhinal cortex atrophy. A voxel-based morphometry study* (Di Paola et al., 2007) | AD, 64 years, n = 18<br><br>HC, 65 years, n = 18 | HC > AD | $p < 0.05$<br><br>Corrected for multiple comparisons | 1.5 T<br><br>Talairach<br><br>SPM2 |
| *Fronto-temporal-lobe atrophy in early-stage Alzheimer's disease identified using an improved detection methodology* (Farrow et al., 2007) | Early stage AD, First scan (AD1), 77 years, n = 7<br><br>HC, 70 years, n = 11 | HC > AD1 | $p < 0.05$<br><br>Corrected for multiple comparisons | 1.5 T<br><br>Talairach<br><br>SPM2 |
| *Detection of grey matter loss in mild Alzheimer's disease with voxel based morphometry* (Frisoni et al., 2002) | AD, 74 years, n = 29<br><br>HC, 74 years, n = 26 | HC > AD | $p < 0.05$<br><br>Corrected for multiple comparisons | 1.5 T<br><br>MNI<br><br>SPM99 |
| *Regional brain atrophy and functional disconnection across Alzheimer's disease evolution* (Gili et al., 2011) | AD, 72 years, n = 11<br><br>HC, 64 years, n = 10 | HC > AD | $p < 0.001$<br><br>Cluster-wise corrected for multiple comparisons | 3.0 T<br><br>MNI<br><br>SPM5 |
| *Correlation between Topographic N400 Anomalies and Reduced Cerebral Blood Flow in the Anterior Temporal Lobes of Patients with Dementia* (Grieder et al., 2013) | AD, 66 years, n = 14<br><br>HC, 69 years, n = 19 | HC > AD | $p < 0.01$<br><br>Voxel-wise corrected for multiple comparisons | 3.0 T<br><br>MNI<br><br>SPM8 |

| | | | | |
|---|---|---|---|---|
| *Voxel-based assessment of gray and white matter volumes in Alzheimer's disease* (Guo et al., 2010) | AD, 72 years, n = 13<br><br>HC, 70 years, n = 14 | HC > AD | $p < 0.05$<br><br>Corrected for multiple comparisons | 3.0 T<br><br>MNI<br><br>SPM2 |
| *Basal forebrain atrophy is a presymptomatic marker for Alzheimer's disease* (Hall et al., 2008) | Probable AD, 83 years, n = 26<br><br>HC, 77 years, n = 127 | HC > Probable AD | $p < 0.05$<br><br>Voxel-wise corrected for multiple comparisons | 1.5 T<br><br>MNI<br><br>SPM2 |
| *Increased fMRI responses during encoding in mild cognitive impairment* (Hämäläinen et al., 2007) | AD, 73 years, n = 15<br><br>HC, 71 years, n = 21 | HC > AD | $p < 0.01$<br><br>Cluster-wise corrected for multiple comparisons | 1.5 T<br><br>MNI<br><br>SPM2 |
| *Cardiorespiratory fitness and preserved medial temporal lobe volume in Alzheimer disease* (Honea et al., 2009) | Early AD, 74 years, n = 60<br><br>HC, 73 years, n = 56 | HC > Early AD | $p < 0.05$<br><br>Corrected for multiple comparisons | 3.0 T<br><br>MNI<br><br>SPM5 |
| *Neural Substrates of Semantic Prospection - Evidence from the Dementias* (Irish et al., 2016) | AD, 65 years, n = 15<br><br>HC, 67 years, n = 20 | HC > AD | $p < 0.05$<br><br>Voxel-wise corrected for multiple comparisons | 3.0 T<br><br>MNI<br><br>FSL |
| *Episodic future thinking is impaired in the behavioral variant of frontotemporal dementia* (Irish et al., 2013) | AD, 65 years, n = 10<br><br>HC, 69 years, n = 10 | HC > AD | $p < 0.001$ | 3.0 T<br><br>MNI |

| | | | | |
|---|---|---|---|---|
| | | | Voxel-wise corrected for multiple comparisons | FSL |
| *Grey and white matter correlates of recent and remote autobiographical memory retrieval-insights from the dementias* (Irish et al., 2014) | AD, 68 years, n = 15<br><br>HC, 72 years, n = 14 | HC > AD | $p < 0.001$<br><br>Voxel-wise corrected for multiple comparisons | 3.0 T<br><br>MNI<br><br>FSL |
| *Comparison of gray matter and metabolic reduction in mild Alzheimer's disease using FDG-PET and voxel-based morphometric MR studies* (Ishii et al., 2005) | Mild AD, 67 years, n = 30<br><br>HC, 67 years, n = 30 | HC > Mild AD | $p < 0.05$<br><br>Voxel-wise corrected for multiple comparisons | 1.5 T<br><br>MNI<br><br>SPM99 |
| *Comparison of grey matter and metabolic reductions in frontotemporal dementia using FDG-PET and voxel-based morphometric MR studies* (Kanda et al., 2008) | AD, 65 years, n = 20<br><br>HC, 65 years, n = 20 | HC > AD | $p < 0.01$<br><br>Voxel-wise corrected for multiple comparisons | 1.5 T<br><br>MNI<br><br>SPM2 |
| *Voxel-based morphometric study of brain volume changes in patients with Alzheimer's disease assessed according to the Clinical Dementia Rating score* (Kim et al., 2011) | AD, 69 years, n = 61<br><br>HC, 70 years, n = 33 | HC > AD | $p < 0.01$<br><br>Corrected for multiple comparisons | 3.0 T<br><br>MNI<br><br>SPM2 |
| *Self-appraisal in behavioral variant frontotemporal degeneration* (Massimo et al., 2013) | AD, 71 years, n = 1773<br><br>HC, 64 years, n = 30 | HC > AD | $p < 0.05$<br><br>Cluster-wise corrected for | Unknown MRI field strength<br><br>Talairach |

| | | | | |
|---|---|---|---|---|
| | | | multiple comparisons | SPM5 |
| *Longitudinal evaluation of both morphologic and functional changes in the same individuals with Alzheimer's disease* (Matsuda et al., 2002) | AD, 71 years, n = 15<br><br>HC, 71 years, n = 25 | HC > AD | $p < 0.05$<br><br>Corrected for multiple comparisons | 1.0 T<br>MNI<br><br>SPM99 |
| *In vivo SPECT imaging of vesicular acetylcholine transporter using [(123)I]-IBVM in early Alzheimer's disease* (Mazère et al., 2008) | AD, 81 years, n = 8<br><br>HC, 74 years, n = 8 | HC > AD patients | $p < 0.01$<br><br>Cluster-wise corrected for multiple comparisons | 1.5 T<br><br>Talairach<br><br>Unknown software system |
| *Changes in brain morphology in Alzheimer disease and normal aging: Is Alzheimer disease an exaggerated aging process?* (Ohnishi et al., 2001) | AD, 72 years, n = 26<br><br>HC, 71 years, n = 22 | HC > AD | $p < 0.001$ | 1.0 T<br><br>MNI<br><br>SPM96 |
| *Distinct MRI atrophy patterns in autopsy-proven Alzheimer's disease and frontotemporal lobar degeneration* (Rabinovici et al., 2007) | AD, 65 years, n = 11<br><br>HC, 64 years, n = 40 | HC > AD | $p < 0.001$ | 1.5 T<br><br>MNI<br><br>SPM2 |
| *Voxel based morphometry features and follow-up of amnestic patients at high risk for Alzheimer's disease conversion* (Rami et al., 2009) | AD, 76 years, n = 31<br><br>HC, 74 years, n = 27 | C > AD | $p < 0.001$ | 1.5 T<br><br>MNI<br><br>SPM2 |

| | | | | |
|---|---|---|---|---|
| *Verbal episodic memory impairment in Alzheimer's disease: a combined structural and functional MRI study* (Rémy et al., 2005) | AD, 72 years, n = 8<br><br>HC, 66 years, n = 11 | HC > AD | $p < 0.001$ | 1.5 T<br><br>MNI<br><br>SPM2 |
| *Four subgroups of Alzheimer's disease based on patterns of atrophy using VBM and a unique pattern for early onset disease* (Shiino et al., 2006) | AD, 71 years, n = 40<br><br>HC, 69 years, n = 88 | HC > AD | $p < 0.05$<br><br>Corrected for multiple comparisons | 1.5 T<br><br>Talairach<br><br>MedX |
| *The Effect of Gray Matter ICA and Coefficient of Variation Mapping of BOLD Data on the Detection of Functional Connectivity Changes in Alzheimer's Disease and bvFTD* (Tuovinen et al., 2016) | AD, 61 years, n = 23<br><br>HC, 59 years, n = 25 | HC > AD | $p < 0.05$<br><br>Voxel-wise corrected for multiple comparisons | 1.5 T<br><br>MNI<br><br>FSL |
| *Disease-specific profiles of apathy in Alzheimer's disease and behavioral-variant frontotemporal dementia differ across the disease course* (Wei et al., 2020) | Early AD, 64 years, n = 10<br><br>HC, 64 years, n = 28 | HC > Early AD | $p < 0.05$<br><br>Voxel-wise corrected for multiple comparisons | 3.0 T<br><br>MNI<br><br>FSL |
| *Imaging correlates of posterior cortical atrophy* (Whitwell et al., 2007) | AD, 65 years, n = 38<br><br>HC, 66 years, n = 38 | HC > AD | $p < 0.05$<br><br>Corrected for multiple comparisons | 1.5 T<br><br>MNI<br><br>SPM2 |
| *Voxel-based detection of white matter abnormalities in mild Alzheimer disease* (Xie et al., 2006) | AD, 72 years, n = 13<br><br>HC, 71 years, n = 16 | HC > AD | $p < 0.001$ | 1.5 T<br><br>MNI |

| | | | | |
|---|---|---|---|---|
| | | | Cluster-wise corrected for multiple comparisons | SPM2 |

**References**

Agosta, F., Pievani, M., Sala, S., Geroldi, C., Galluzzi, S., Frisoni, G., Filippi, M., 2011. White Matter Damage in Alzheimer Disease and Its Relationship to Gray Matter Atrophy. Radiology 258, 853–63. https://doi.org/10.1148/radiol.10101284

Bai, F., Zhang, Z., Yu, H., Shi, Y., Yuan, Y., Zhu, W., Zhang, X., Qian, Y., 2008. Default-mode network activity distinguishes amnestic type mild cognitive impairment from healthy aging: A combined structural and resting-state functional MRI study. Neurosci. Lett. 438, 111–115. https://doi.org/10.1016/j.neulet.2008.04.021

Barbeau, E.J., Ranjeva, J.P., Didic, M., Confort-Gouny, S., Felician, O., Soulier, E., Cozzone, P.J., Ceccaldi, M., Poncet, M., 2008. Profile of memory impairment and gray matter loss in amnestic mild cognitive impairment. Neuropsychologia 46, 1009–1019. https://doi.org/10.1016/j.neuropsychologia.2007.11.019

Baron, J.C., Chételat, G., Desgranges, B., Perchey, G., Landeau, B., de la Sayette, V., Eustache, F., 2001. In vivo mapping of gray matter loss with voxel-based morphometry in mild Alzheimer's disease. NeuroImage 14, 298–309. https://doi.org/10.1006/nimg.2001.0848

Baxter, L.C., Sparks, D.L., Johnson, S.C., Lenoski, B., Lopez, J.E., Connor, D.J., Sabbagh, M.N., 2006. Relationship of cognitive measures and gray and white matter in Alzheimer's disease. J. Alzheimers Dis. JAD 9, 253–260. https://doi.org/10.3233/jad-2006-9304

Bell-McGinty, S., Lopez, O.L., Meltzer, C.C., Scanlon, J.M., Whyte, E.M., Dekosky, S.T., Becker, J.T., 2005. Differential cortical atrophy in subgroups of mild cognitive impairment. Arch. Neurol. 62, 1393–1397. https://doi.org/10.1001/archneur.62.9.1393

Boxer, A.L., Rankin, K.P., Miller, B.L., Schuff, N., Weiner, M., Gorno-Tempini, M.-L., Rosen, H.J., 2003. Cinguloparietal atrophy distinguishes Alzheimer disease from semantic dementia. Arch. Neurol. 60, 949–956. https://doi.org/10.1001/archneur.60.7.949

Bozzali, M., Filippi, M., Magnani, G., Cercignani, M., Franceschi, M., Schiatti, E., Castiglioni, S., Mossini, R., Falautano, M., Scotti, G., Comi, G., Falini, A., 2006. The contribution of voxel-based morphometry in staging patients with mild cognitive impairment. Neurology 67, 453–460. https://doi.org/10.1212/01.wnl.0000228243.56665.c2

Bozzali, M., Giulietti, G., Basile, B., Serra, L., Spanò, B., Perri, R., Giubilei, F., Marra, C., Caltagirone, C., Cercignani, M., 2012. Damage to the cingulum contributes to Alzheimer's disease pathophysiology by deafferentation mechanism. Hum. Brain Mapp. 33, 1295–1308. https://doi.org/10.1002/hbm.21287

Brenneis, C., Wenning, G., Egger, K., Schocke, M., Trieb, T., Seppi, K., Marksteiner, J., Ransmayr, G., Benke, T., Poewe, W., 2004. Basal forebrain atrophy is a distinctive pattern in dementia with Lewy bodies. Neuroreport 15, 1711–4. https://doi.org/10.1097/01.wnr.0000136736.73895.03

Caroli, A., Testa, C., Geroldi, C., Nobili, F., Barnden, L.R., Guerra, U.P., Bonetti, M., Frisoni, G.B., 2007. Cerebral perfusion correlates of conversion to Alzheimer's disease in amnestic mild cognitive impairment. J. Neurol. 254, 1698–1707. https://doi.org/10.1007/s00415-007-0631-7

Chételat, G., Desgranges, B., Sayette, V., Viader, F., Eustache, F., Baron, J.-C., 2002. Mapping gray matter loss with voxel-based morphometry in mild cognitive impairment. Neuroreport 13, 1939–43. https://doi.org/10.1097/00001756-200210280-00022

Colloby, S.J., O'Brien, J.T., Taylor, J.-P., 2014. Patterns of cerebellar volume loss in dementia with Lewy bodies and Alzheimer's disease: A VBM-DARTEL study. Psychiatry Res. 223, 187–191. https://doi.org/10.1016/j.pscychresns.2014.06.006

Dashjamts, T., Yoshiura, T., Hiwatashi, A., Yamashita, K., Monji, A., Ohyagi, Y., Kamano, H., Kawashima, T., Kira, J.-I., Honda, H., 2011. Simultaneous arterial spin labeling cerebral blood flow and morphological assessments for detection of Alzheimer's disease. Acad. Radiol. 18, 1492–1499. https://doi.org/10.1016/j.acra.2011.07.015

Dermody, N., Hornberger, M., Piguet, O., Hodges, J.R., Irish, M., 2016a. Prospective Memory Impairments in Alzheimer's Disease and Behavioral Variant Frontotemporal Dementia: Clinical and Neural Correlates. J. Alzheimers Dis. JAD 50, 425–441. https://doi.org/10.3233/JAD-150871

Dermody, N., Wong, S., Ahmed, R., Piguet, O., Hodges, J.R., Irish, M., 2016b. Uncovering the Neural Bases of Cognitive and Affective Empathy Deficits in Alzheimer's Disease and the Behavioral-Variant of Frontotemporal Dementia. J. Alzheimers Dis. JAD 53, 801–816. https://doi.org/10.3233/JAD-160175

Di Paola, M., Macaluso, E., Carlesimo, G.A., Tomaiuolo, F., Worsley, K.J., Fadda, L., Caltagirone, C., 2007. Episodic memory impairment in patients with Alzheimer's disease is correlated with entorhinal cortex atrophy. A voxel-based morphometry study. J. Neurol. 254, 774–781. https://doi.org/10.1007/s00415-006-0435-1

Farrow, T.F.D., Thiyagesh, S.N., Wilkinson, I.D., Parks, R.W., Ingram, L., Woodruff, P.W.R., 2007. Fronto-temporal-lobe atrophy in early-stage Alzheimer's disease identified using an improved detection methodology. Psychiatry Res. 155, 11–19. https://doi.org/10.1016/j.pscychresns.2006.12.013

Frisoni, G.B., Testa, C., Zorzan, A., Sabattoli, F., Beltramello, A., Soininen, H., Laakso, M.P., 2002. Detection of grey matter loss in mild Alzheimer's disease with voxel based morphometry. J. Neurol. Neurosurg. Psychiatry 73, 657–664. https://doi.org/10.1136/jnnp.73.6.657

Gili, T., Cercignani, M., Serra, L., Perri, R., Giove, F., Maraviglia, B., Caltagirone, C., Bozzali, M., 2011. Regional brain atrophy and functional disconnection across Alzheimer's disease evolution. J. Neurol. Neurosurg. Psychiatry 82, 58–66. https://doi.org/10.1136/jnnp.2009.199935

Gold, B.T., Jiang, Y., Jicha, G.A., Smith, C.D., 2010. Functional response in ventral temporal cortex differentiates mild cognitive impairment from normal aging. Hum. Brain Mapp. 31, 1249–1259. https://doi.org/10.1002/hbm.20932

Grieder, M., Crinelli, R.M., Jann, K., Federspiel, A., Wirth, M., Koenig, T., Stein, M., Wahlund, L.-O., Dierks, T., 2013. Correlation between Topographic N400 Anomalies and Reduced Cerebral Blood Flow in the Anterior Temporal Lobes of Patients with Dementia. J. Alzheimers Dis. JAD 36, 711–731. https://doi.org/10.3233/JAD-121690

Guo, X., Wang, Z., Li, K., Li, Z., Qi, Z., Jin, Z., Yao, L., Chen, K., 2010. Voxel-based assessment of gray and white matter volumes in Alzheimer's disease. Neurosci. Lett. 468, 146–150. https://doi.org/10.1016/j.neulet.2009.10.086

Hall, A.M., Moore, R.Y., Lopez, O.L., Kuller, L., Becker, J.T., 2008. Basal forebrain atrophy is a presymptomatic marker for Alzheimer's disease. Alzheimers Dement. J. Alzheimers Assoc. 4, 271–279. https://doi.org/10.1016/j.jalz.2008.04.005

Hämäläinen, A., Pihlajamäki, M., Tanila, H., Hänninen, T., Niskanen, E., Tervo, S., Karjalainen, P.A., Vanninen, R.L., Soininen, H., 2007. Increased fMRI responses during encoding in mild cognitive impairment. Neurobiol. Aging 28, 1889–1903. https://doi.org/10.1016/j.neurobiolaging.2006.08.008

Honea, R.A., Thomas, G.P., Harsha, A., Anderson, H.S., Donnelly, J.E., Brooks, W.M., Burns, J.M., 2009. Cardiorespiratory fitness and preserved medial temporal lobe volume in Alzheimer disease. Alzheimer Dis. Assoc. Disord. 23, 188–197. https://doi.org/10.1097/WAD.0b013e31819cb8a2

Irish, M., Eyre, N., Dermody, N., O'Callaghan, C., Hodges, J.R., Hornberger, M., Piguet, O., 2016. Neural Substrates of Semantic Prospection - Evidence from the Dementias. Front. Behav. Neurosci. 10, 96. https://doi.org/10.3389/fnbeh.2016.00096

Irish, M., Hodges, J.R., Piguet, O., 2013. Episodic future thinking is impaired in the behavioural variant of frontotemporal dementia. Cortex J. Devoted Study Nerv. Syst. Behav. 49, 2377–2388. https://doi.org/10.1016/j.cortex.2013.03.002

Irish, M., Hornberger, M., El Wahsh, S., Lam, B.Y.K., Lah, S., Miller, L., Hsieh, S., Hodges, J.R., Piguet, O., 2014. Grey and white matter correlates of recent and remote autobiographical memory retrieval-insights from the dementias. PloS One 9, e113081. https://doi.org/10.1371/journal.pone.0113081

Ishii, K., Sasaki, H., Kono, A.K., Miyamoto, N., Fukuda, T., Mori, E., 2005. Comparison of gray matter and metabolic reduction in mild Alzheimer's disease using FDG-PET and voxel-based morphometric MR studies. Eur. J. Nucl. Med. Mol. Imaging 32, 959–963. https://doi.org/10.1007/s00259-004-1740-5

Kanda, T., Ishii, K., Uemura, T., Miyamoto, N., Yoshikawa, T., Kono, A.K., Mori, E., 2008. Comparison of grey matter and metabolic reductions in frontotemporal dementia using FDG-PET and voxel-based morphometric MR studies. Eur. J. Nucl. Med. Mol. Imaging 35, 2227–2234. https://doi.org/10.1007/s00259-008-0871-5

Kim, S., Youn, Y.C., Hsiung, G.-Y.R., Ha, S.-Y., Park, K.-Y., Shin, H.-W., Kim, D.-K., Kim, S.-S., Kee, B.S., 2011. Voxel-based morphometric study of brain volume changes in patients with Alzheimer's disease assessed according to the Clinical Dementia Rating score. J. Clin. Neurosci. Off. J. Neurosurg. Soc. Australas. 18, 916–921. https://doi.org/10.1016/j.jocn.2010.12.019

Long, Z., Jing, B., Yan, H., Dong, J., Liu, H., Mo, X., Han, Y., Li, H., 2016. A support vector machine-based method to identify mild cognitive impairment with multi-level characteristics of magnetic resonance imaging. Neuroscience 331, 169–176. https://doi.org/10.1016/j.neuroscience.2016.06.025

Massimo, L., Libon, D.J., Chandrasekaran, K., Dreyfuss, M., McMillan, C.T., Rascovsky, K., Boller, A., Grossman, M., 2013. Self-appraisal in behavioural variant frontotemporal degeneration. J. Neurol. Neurosurg. Psychiatry 84, 148–153. https://doi.org/10.1136/jnnp-2012-303153

Matsuda, H., Kitayama, N., Ohnishi, T., Asada, T., Nakano, S., Sakamoto, S., Imabayashi, E., Katoh, A., 2002. Longitudinal evaluation of both morphologic and functional changes in the same individuals with Alzheimer's disease. J. Nucl. Med. Off. Publ. Soc. Nucl. Med. 43, 304–311.

Mazère, J., Prunier, C., Barret, O., Guyot, M., Hommet, C., Guilloteau, D., Dartigues, J.F., Auriacombe, S., Fabrigoule, C., Allard, M., 2008. In vivo SPECT imaging of vesicular acetylcholine transporter using [(123)I]-IBVM in early Alzheimer's disease. NeuroImage 40, 280–288. https://doi.org/10.1016/j.neuroimage.2007.11.028

Müller, Cieslik, E.C., Laird, A.R., Fox, P.T., Radua, J., Mataix-Cols, D., Tench, C.R., Yarkoni, T., Nichols, T.E., Turkeltaub, P.E., Wager, T.D., Eickhoff, S.B., 2018. Ten simple rules for neuroimaging meta-analysis. Neurosci. Biobehav. Rev. 84, 151–161. https://doi.org/10.1016/j.neubiorev.2017.11.012

Ohnishi, T., Matsuda, H., Tabira, T., Asada, T., Uno, M., 2001. Changes in brain morphology in Alzheimer disease and normal aging: is Alzheimer disease an exaggerated aging process? AJNR Am. J. Neuroradiol. 22, 1680–1685.

Pennanen, C., Testa, C., Laakso, M.P., Hallikainen, M., Helkala, E.-L., Hänninen, T., Kivipelto, M., Könönen, M., Nissinen, A., Tervo, S., Vanhanen, M., Vanninen, R., Frisoni, G.B., Soininen, H.,

2005. A voxel based morphometry study on mild cognitive impairment. J. Neurol. Neurosurg. Psychiatry 76, 11–14. https://doi.org/10.1136/jnnp.2004.035600

Rabinovici, G.D., Seeley, W.W., Kim, E.J., Gorno-Tempini, M.L., Rascovsky, K., Pagliaro, T.A., Allison, S.C., Halabi, C., Kramer, J.H., Johnson, J.K., Weiner, M.W., Forman, M.S., Trojanowski, J.Q., Dearmond, S.J., Miller, B.L., Rosen, H.J., 2007. Distinct MRI atrophy patterns in autopsy-proven Alzheimer's disease and frontotemporal lobar degeneration. Am. J. Alzheimers Dis. Other Demen. 22, 474–488. https://doi.org/10.1177/1533317507308779

Rami, L., Gómez-Anson, B., Monte, G.C., Bosch, B., Sánchez-Valle, R., Molinuevo, J.L., 2009. Voxel based morphometry features and follow-up of amnestic patients at high risk for Alzheimer's disease conversion. Int. J. Geriatr. Psychiatry 24, 875–884. https://doi.org/10.1002/gps.2216

Rémy, F., Mirrashed, F., Campbell, B., Richter, W., 2005. Verbal episodic memory impairment in Alzheimer's disease: a combined structural and functional MRI study. NeuroImage 25, 253–266. https://doi.org/10.1016/j.neuroimage.2004.10.045

Schmidt-Wilcke, T., Poljansky, S., Hierlmeier, S., Hausner, J., Ibach, B., 2009. Memory performance correlates with gray matter density in the ento-/perirhinal cortex and posterior hippocampus in patients with mild cognitive impairment and healthy controls — A voxel based morphometry study. NeuroImage 47, 1914–1920. https://doi.org/10.1016/j.neuroimage.2009.04.092

Shiino, A., Watanabe, T., Maeda, K., Kotani, E., Akiguchi, I., Matsuda, M., 2006. Four subgroups of Alzheimer's disease based on patterns of atrophy using VBM and a unique pattern for early-onset disease. NeuroImage 33, 17–26. https://doi.org/10.1016/j.neuroimage.2006.06.010

Tuovinen, T., Rytty, R., Moilanen, V., Abou Elseoud, A., Veijola, J., Remes, A.M., Kiviniemi, V.J., 2016. The Effect of Gray Matter ICA and Coefficient of Variation Mapping of BOLD Data on the Detection of Functional Connectivity Changes in Alzheimer's Disease and bvFTD. Front. Hum. Neurosci. 10, 680. https://doi.org/10.3389/fnhum.2016.00680

Wei, G., Irish, M., Hodges, J.R., Piguet, O., Kumfor, F., 2020. Disease-specific profiles of apathy in Alzheimer's disease and behavioural-variant frontotemporal dementia differ across the disease course. J. Neurol. 267, 1086–1096. https://doi.org/10.1007/s00415-019-09679-1

Whitwell, J.L., Jack, C.R., Kantarci, K., Weigand, S.D., Boeve, B.F., Knopman, D.S., Drubach, D.A., Tang-Wai, D.F., Petersen, R.C., Josephs, K.A., 2007. Imaging correlates of posterior cortical atrophy. Neurobiol. Aging 28, 1051–1061. https://doi.org/10.1016/j.neurobiolaging.2006.05.026

Xie, S., Xiao, J.X., Gong, G.L., Zang, Y.F., Wang, Y.H., Wu, H.K., Jiang, X.X., 2006. Voxel-based detection of white matter abnormalities in mild Alzheimer disease. Neurology 66, 1845–1849. https://doi.org/10.1212/01.wnl.0000219625.77625.aa

Yin, C., Yi, L., Jia, L., Wang, J., Liu, P., Guo, Y., Han, Y., 2014. Early morphological brain abnormalities in patients with amnestic mild cognitive impairment. Transl. Neurosci. 5, 253–259. https://doi.org/10.2478/s13380-014-0234-6

Zhao, Z., Lu, J., Jia, X., Chao, W., Han, Y., Jia, J., Li, K., 2014. Selective Changes of Resting-State Brain Oscillations in aMCI: An fMRI Study Using ALFF. BioMed Res. Int. 2014, 1–7. https://doi.org/10.1155/2014/920902
